# Metabolic trade-offs in response to increased acetate metabolism in *Escherichia coli*

**DOI:** 10.1101/2024.11.20.624476

**Authors:** Jennifer T. Pentz, Samuel I. Koehler, Earl A. Middlebrook, Kayley T. You Mak, Blake T. Hovde, Erik R. Hanschen

## Abstract

Adaptive laboratory evolution (ALE) is a powerful tool to improve phenotypes in microbial cell factories for biotechnological processes. For example, ALE can be utilized to improve growth on alternative carbon sources such as acetate. Acetate is becoming a promising alternative feedstock to produce a variety of bio-products, but growth on acetate is slow relative to other preferred carbon sources and can be toxic to cells. However, ALE in a homogenous environment can lead to evolutionary trade-offs for growth on other carbon sources, which might be detrimental for feedstocks that contain multiple sugars. Here, we evolved *Escherichia coli* for ∼100 generations using continuous culturing in turbidostats in media containing acetate as the sole carbon source to rapidly select for increased growth in acetate. We measured absolute fitness of 119 clones in acetate and glucose separately to characterize trade-offs and found that trade-off patterns were heterogenous amongst measured clones, with 45% of clones showing significantly reduced growth in glucose. Sequencing revealed that the trade-offs were a result of antagonistic pleiotropic effects of early adaptive mutations, but not all high-effect mutations conferred a trade-off, consistent with the model that mutations accumulated in the selected environment can display a range of pleiotropic effects in other environments. Together, these results suggest that ALE in a homogeneous environment needs to be carefully considered for the improvement of metabolic traits in microbial cells as this could limit the use of evolved strains in complex feedstocks that contain multiple sugars.

## Introduction

During adaptive evolution, novel mutations with increased or neutral fitness in one environment can reduce it in another^1,2^. Trade-offs may emerge due to selective or neutral processes. Selection may favor beneficial mutations in the current environment that are deleterious in other environments, termed antagonistic pleiotropy^3,4^. Alternatively, neutral mutations that accumulate in the selected environment can result in reduced fitness in non-selected environments, also referred to as mutation accumulation. Characterizing trade-offs are of great interest because these may constrain evolutionary trajectories^1,2^ as well as influence the ‘evolvability’, the ability of an organism to generate beneficial mutations, of a system^5–7^.

Understanding the emergence of trade-offs when adapting organisms to high density or in novel culture conditions is critical for metabolic engineering efforts to produce desired products. Microbial evolution experiments have been a powerful approach for demonstrating and characterizing the existence of such trade-offs^3,8–15^. Furthermore, directed evolution is increasingly being used in industrially relevant microorganisms for new and/or improved functions, such as industrial enzyme optimization^16,17^. One common utility for directed evolution in industry is to maximize carbon source utilization^18,19^ despite the evolutionary trade-offs that might arise. Understanding these emergent trade-offs is critical for optimizing metabolic engineering strategies and achieving desired industrial outcomes with adaptive laboratory evolution (ALE).

Acetate is a promising alternative carbon source to produce a variety of desired bioproducts^20,21^. Furthermore, acetate can improve the economic feasibility of various bioprocesses as it is relatively inexpensive^22^, is present at high levels in waste streams^23,24^, and is generated as a byproduct during the pretreatment of lignocellulosic biomass^25–27^. *Escherichia coli* produces acetate as a by-product of glycolytic metabolism, or overflow metabolism. Historically, acetate accumulation during fermentation was though to inhibit microbial growth, motivating many efforts in *E. coli* to limit acetate accumulation or increase the tolerance of cells to the presence of acetate^28–31^. *E. coli*, however, can also utilize acetate as an alternative carbon source ^32^ and has been explored for its ability to produce a variety of biochemicals from acetate (see Kutscha and Pflügl 2020 and references within). *E. coli* assimilate acetate by converting it to acetyl-CoA either through the Pta-AckA pathway or the acetyl-CoA synthase (Acs) that can then generate energy via the tricarboxylic acid (TCA) cycle^33–35^. However, the growth and consumption rates of *E. coli* on acetate are much lower when compared to preferred carbon sources such as glucose^28,29,36^. Therefore, ALE could be a promising approach to increase the rate of acetate utilization as well as improve tolerance to the toxic effects that have been documented^28–30,36^. Indeed, there have been a few examples of increased acetate consumption or tolerance in response to directed evolution in *E. coli*^32,37,38^, but there has been little investigation into the emergent metabolic trade-offs that could, for example, affect the potential for co-substrate utilization with glycolytic nutrients. Understanding these trade-offs will be important when adapting organisms to better utilize alternative carbon sources such as acetate, a key challenge in optimizing industrial processes.

Here we utilized experimental evolution in long-term culturing devices, or chemostats, with *E. coli* to select for increased efficiency of acetate metabolism. We chose *E. coli* because it is a widely used production species and there is a large focus for strain development for increased acetate metabolism and tolerance. We then examined the emergence of trade-offs for growth in a preferred carbon source, glucose. After ∼100 generations of chemostat evolution, evolved populations and isolated clones therein grew significantly faster than the wild-type *E. coli* BW25113 ancestor in media containing acetate as the sole carbon source. However, some of the evolved isolates displayed fitness trade-offs when grown in glucose media whereas others did not show a measurable trade-off. This was likely due to the underlying genetics of increased acetate metabolism and tolerance as observed trade-offs in glucose correlated with different genotypes isolated from evolved chemostat populations. Overall, our results suggest that directed evolution for increased acetate metabolism rapidly (less than 100 generations) results in trade-offs for glucose metabolism due to the antagonistic pleiotropic effects of adaptive mutations, which should be considered when utilizing artificial laboratory evolution in homogenous environments for evolving strains for industrial processes involving complex environments.

## Results

### Rapid evolution of increased growth in media with acetate as the sole carbon source

*E. coli* can consume acetate as a sole carbon source, but it is less preferred and can even inhibit bacterial growth^28–30^. To select for increased metabolism of acetate in *E. coli* BW25113, we utilized adaptive laboratory evolution (ALE) using chemostats, or continuous culturing devices^39^. Key advantages of using chemostats for ALE are bacteria can be grown in a defined and constant environment, the growth rate of cells can be controlled, and the effective population size of evolving cultures is large allowing for rapid exploration of genotype and phenotype space^40–42^. We used Chi.Bio bioreactors for our ALE, an all-in-one continuous culturing system that includes a control computer, individual reactors, and peristaltic pumps for the inflow of fresh media [minimal media (MM) with 17.3 mM acetate as the sole carbon source, see Methods] and the outflow of waste (Figure 1A). We operated the reactors in turbidostat mode, which maintained cultures within 2% of the optical density (OD) set point of 1.0, corresponding to roughly 4 × 10^9^ cells/mL (Figure 1, Figure S1). We evolved six independent cultures originating from a single clone of *E. coli* BW25113 in this environment for ∼12.5 days, resulting in ∼90 generations of bacterial evolution (Figure 1, Figure S1,S2).

**Figure 1.**
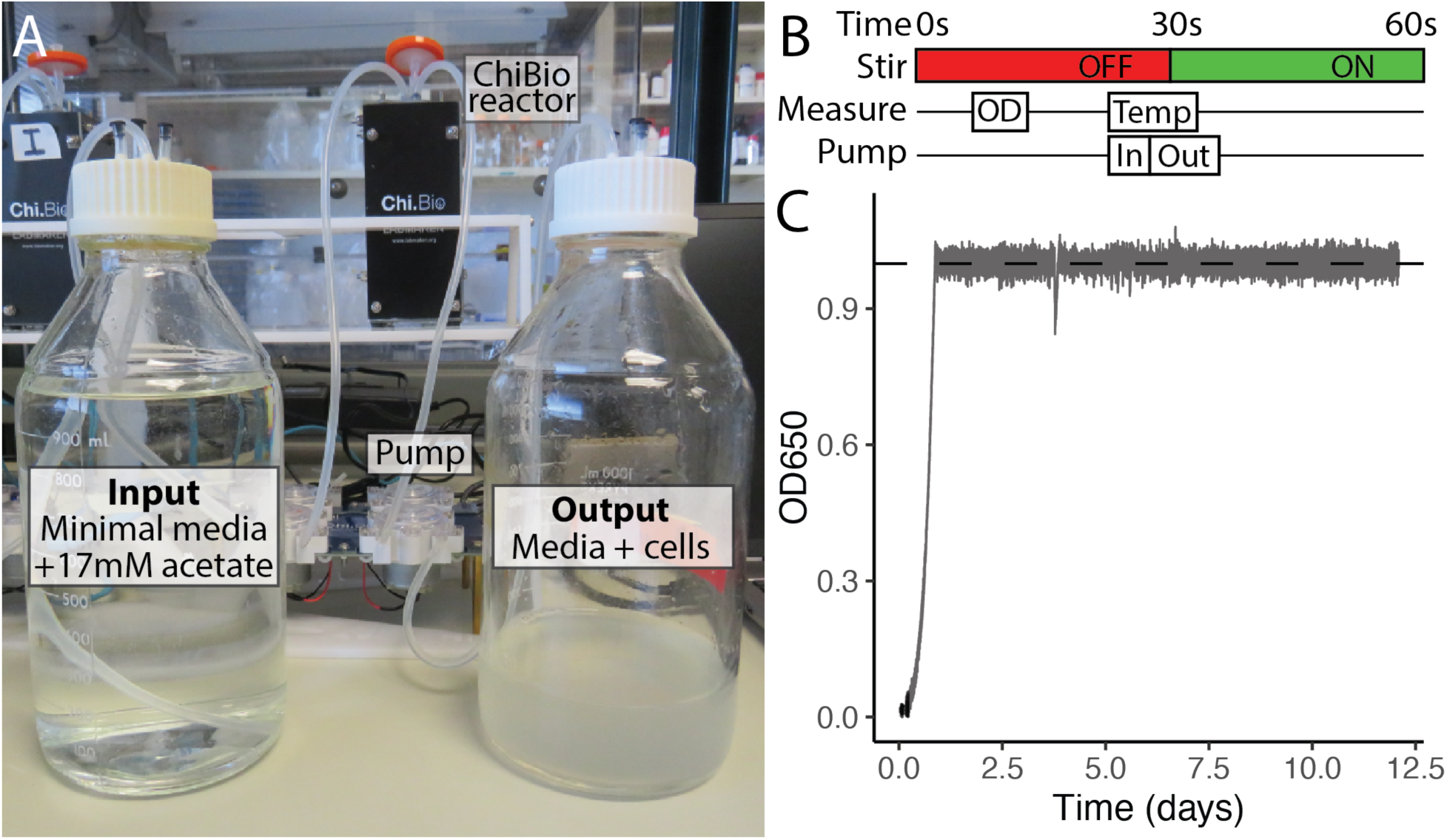
Chi.Bio chemostat automated culturing platform and the evolution of increased acetate metabolism. A) We used Chi.Bio reactors to select for increased acetate metabolism in *E. coli*. Shown is the Chi.Bio reactor on top of a custom laser-cut holding stand and the Chi.Bio peristaltic pump. The input fresh media was minimal media with 17.3mM acetate as the sole carbon source. B) Schematic of 60-second duty cycle used for our adaptive laboratory evolution (ALE, adapted from Steel et al. 2020). Timeline ‘Stir’ indicate the state of the stir bar as either stationary (OFF) or spinning (ON), ‘Measure’ identifies when optical density (OD, measured at 650 nm) and temperature (Temp) measurements are taken, and ‘Pump’ indicates when fresh media is being pumped into the system (In) and waste media is being pumped out (Out). C) Experimental raw data from one of six chemostats (Chemostat A) over the 12.5-day experiment. All chemostats were run in turbidostat mode with an OD set-point of 1.0 See also Figure S1, S2.

We next determined if end-point populations grew faster in acetate than their wild-type (WT) ancestor. Using the Chi.Bio chemostats, we measured the growth rate of the evolved populations as well as the ancestor with the OD set to follow a dithered waveform, which rapidly dilutes the culture to a lower OD once the user-defined upper limit is reached (Figure 2A; Figure S3), allowing for the precise measurement of growth rate^39^. We developed a custom R script to calculate the slope in between dilutions by identifying the min and max of each cycle and fitting a linear model (Figure 2A)^43^. All evolved populations grew ∼2.5 times faster than the ancestor (one-way ANOVA, F_6,282_=98.2, *p*<0.0001, pair-wise differences assessed with Tukey HSD), indicating rapid evolution of increased acetate metabolism.

**Figure 2.**
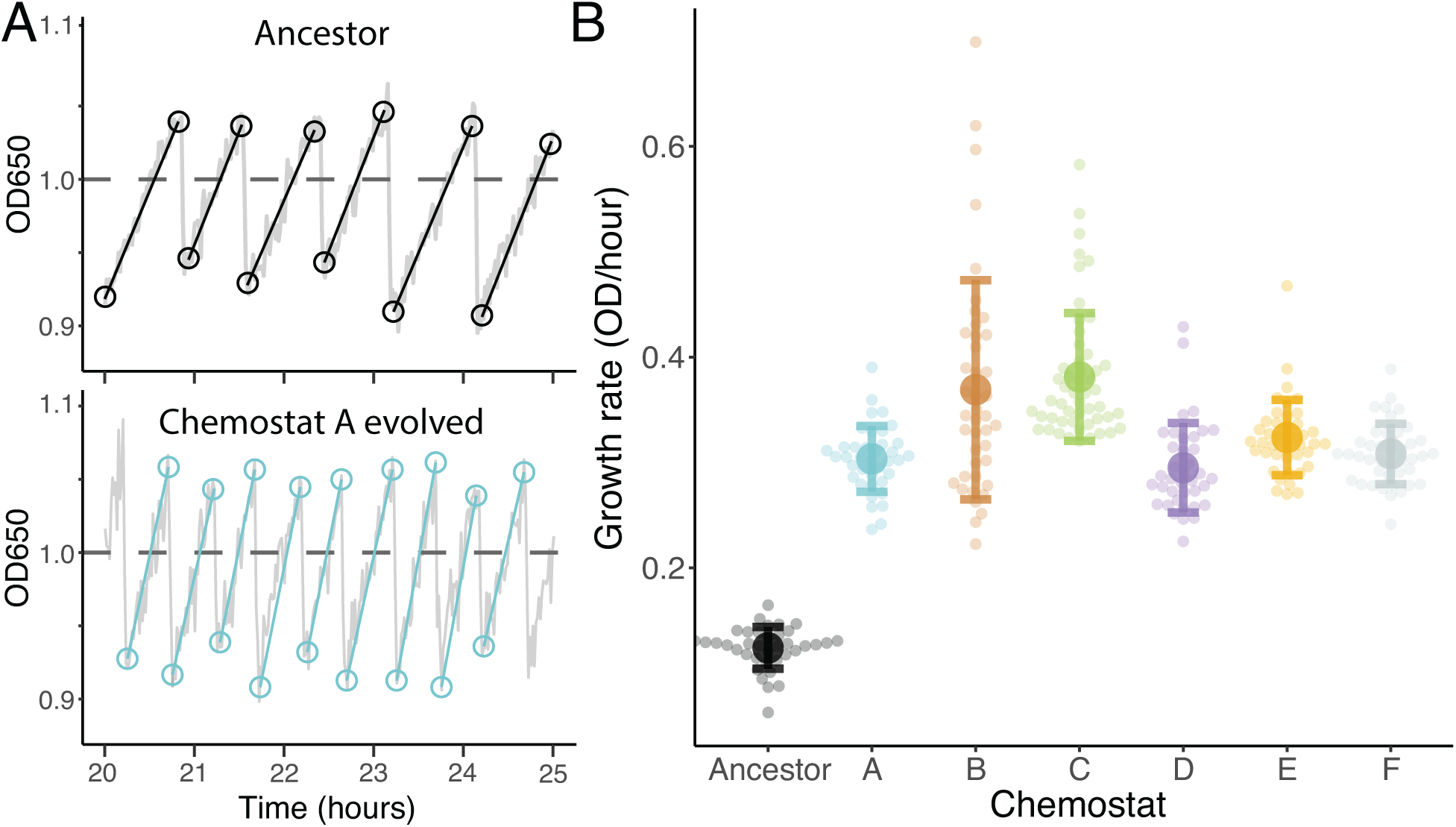
Populations evolved in chemostats show rapid evolution of increased growth in acetate. A) We measured the growth rate of the ancestor and evolved populations using Chi.Bio reactors with the OD (650 nm) set to a dithered waveform with an OD set-point of 1. We calculated the slope between dilutions with a custom R script that automatically finds the max and min of each cycle (circles), then fits a linear model of OD in response to time (see Methods). Figure shows a representative 5 hours of a 20-hour long experiment. B) Evolved populations exhibited an average ∼2.5-fold increase in growth in acetate. Small circles represent individual calculated slopes over 20 hours and large circles represent the average. Error bars show the standard deviation. See also Figure S3.

### Evolved chemostat populations display different trade-off patterns

*E. coli* excrete acetate during exponential growth on glucose, referred to as overflow metabolism, through the Pta-AckA pathway. During stationary phase, however, *E. coli* can uptake and metabolize acetate by two alternative pathways: through the reversible Pta-AckA pathway or via the irreversible acetyl-CoA synthase (Acs), both resulting in acetyl-CoA formation^28^. Therefore, we aimed to determine if clones evolved in an environment with acetate as the sole carbon source exhibited trade-offs when grown in glucose, their preferred carbon source. To do this, we examined the growth of 20 clones isolated from each of the six evolved chemostat populations (19 clones from chemostat C), totaling 119 individual clones, in medium containing either acetate or glucose as the sole carbon source. Specifically, we calculated the maximum growth rate (µ_max_) from growth curves of evolved clones and the ancestor in media with either acetate or glucose measured every 20 minutes in a 96-well plate reader. In acetate, many clones exhibited a significantly higher µ_max_ relative to the ancestor, consistent with population measurements (Figure 3, source data and statistics in Table S1). Surprisingly, many clones from Chemostat C did not show an increase in growth in acetate, with some displaying significantly decreased growth relative to the ancestor which will be further discussed later. Growth trade-offs in glucose were dependent on the chemostat of origin. Many clones grew significantly slower than the ancestor in glucose, indicating a trade-off between acetate and glucose metabolism in these populations. However, other clones showed no measurable trade-off (Figure 3, Table S1), indicating a trade-off between acetate and glucose metabolism is not always inevitable.

**Figure 3.**
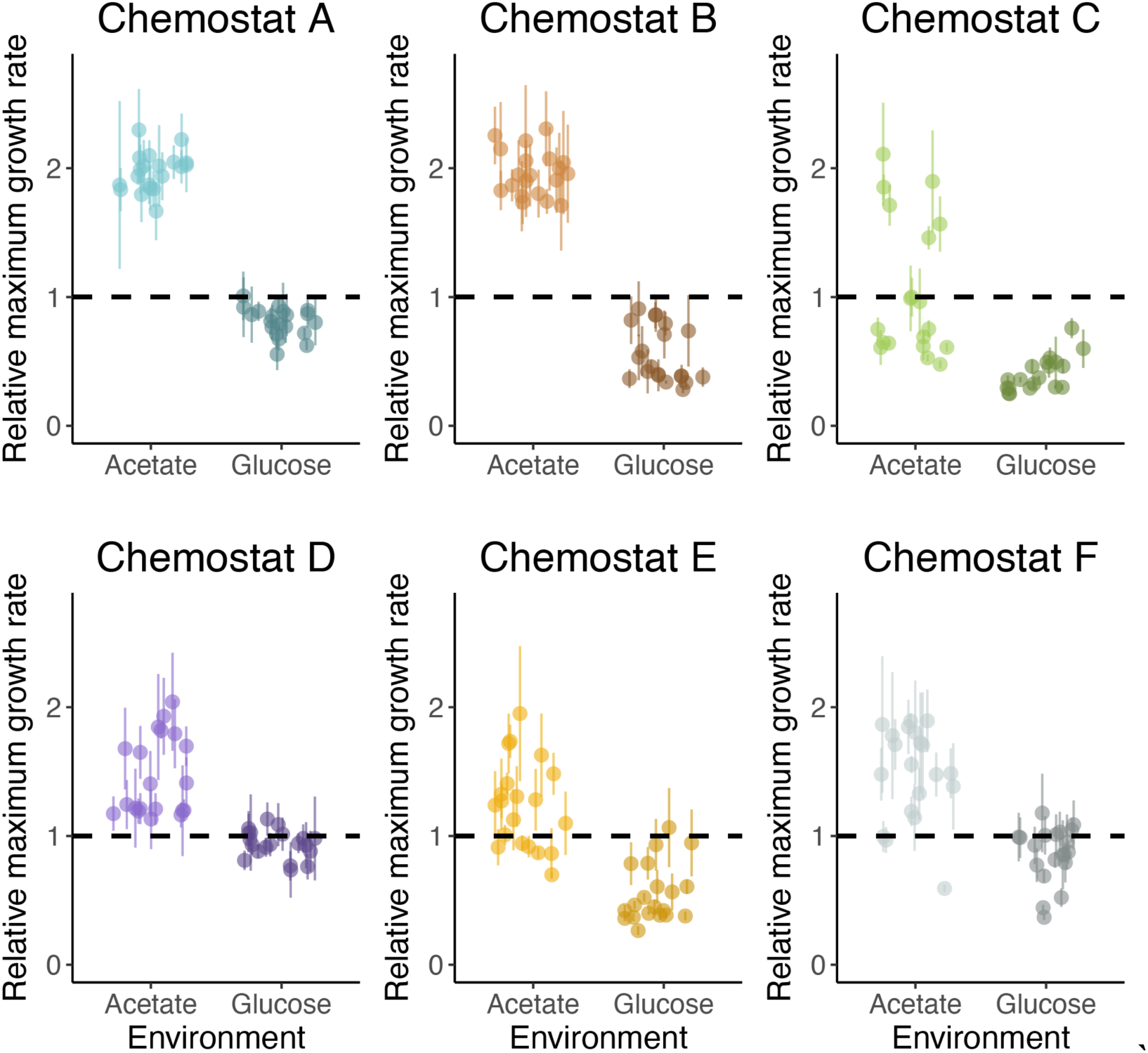
Growth rates of evolved clones relative to the ancestor in acetate or glucose. We isolated 20 clones from each evolved population and calculated their maximum growth rate during exponential growth with a time-resolved 96-well plate reader. Most evolved isolates had increased growth in acetate relative to the ancestor. However, there was variation in the relative growth of evolved isolates in glucose based on the chemostat of origin, suggesting divergence in trade-offs with glucose metabolism associated with increased acetate metabolism. Points represent the mean of four technical replicates for each evolved biological isolate and error bars show the standard deviation. See also Figure S4.

### The genetic basis for increased growth in acetate

To identify the genetic basis of acetate adaptation and observed trade-offs, we sequenced the genomes of all 119 single-strain clones. Different sets of clones isolated from the same chemostat exhibited identical sets of mutations (Figure 4A; clone number listed on top of genotype, mutations listed in Table S2), indicating the presence of a co-existing lineages. On average, most clones had 1-2 mutations apart from chemostat A (∼5 mutations per clone; Figure S5). Most mutations were non-synonymous (roughly 65% missense mutations, 20% nonsense and indel mutations, and 13% synonymous and noncoding mutations), suggesting strong selection (Figure S5). Furthermore, our experimental system had a large population size (∼4×10^9^ cells/mL) implying that drift and hitchhiking are unlikely to have played a role in evolutionary dynamics. Thus, we expect that most mutations that arose had a beneficial fitness effect.

**Figure 4.**
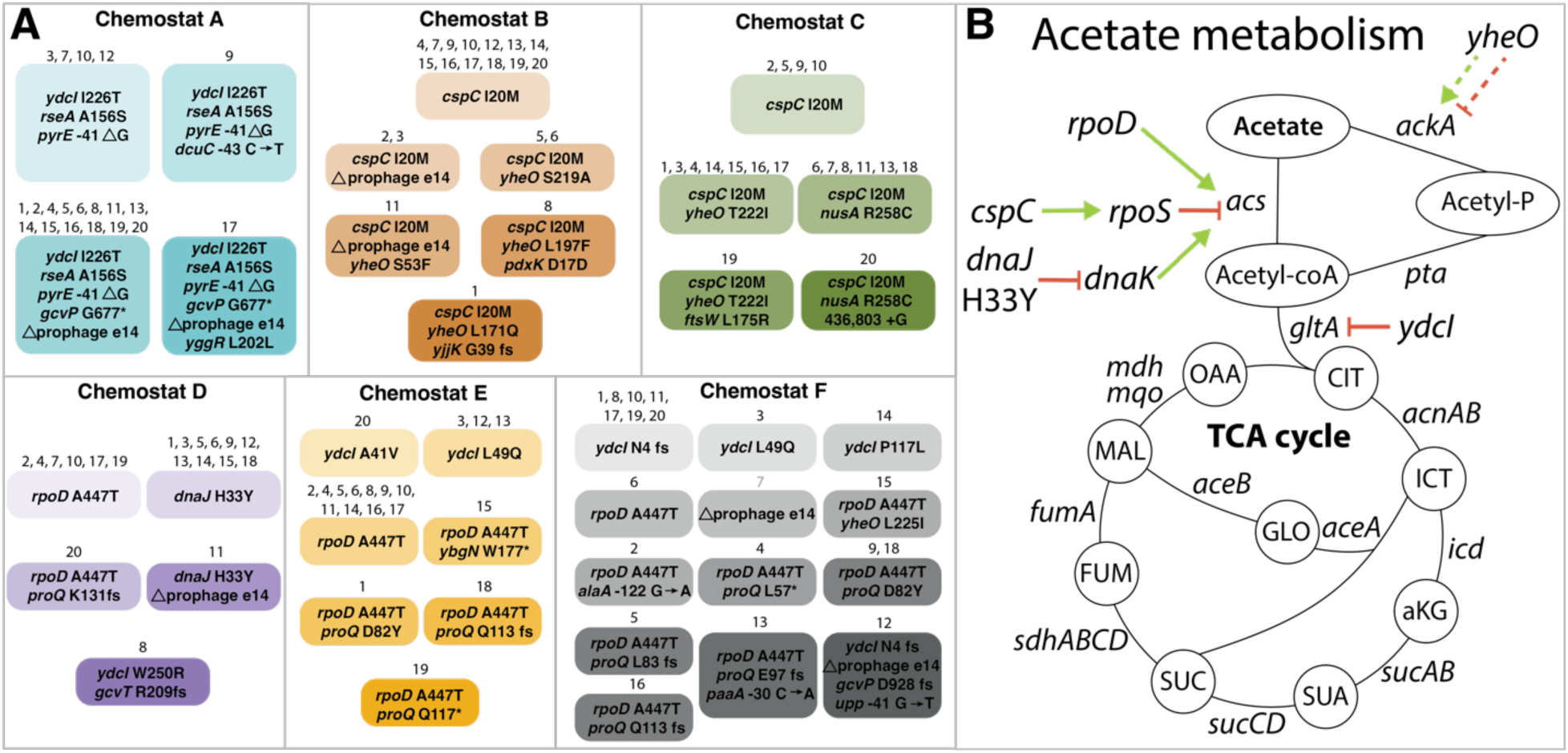
Mutations identified in evolved clones. We sequenced 119 clones isolated from the evolved chemostat populations. Genotypes are shown in bubbles and clone number is designated above. Overall, there were forty genotypes isolated from the six chemostat populations. There was several examples of parallelism across chemostat populations, with mutations occurring in the same gene or even at the nucleotide level [*e.g.*, *cspC* (B, C), *rpoD* (D, E, F), *yheO* (B, C, F), *ydcI* (A, D, E, F), *proQ* (D, E, F), and Δprophage e14 (A, B, C, D)]. Many mutations were predicted to increase the efficiency of acetate metabolism, shown on the diagram of acetate metabolism. B) Simple schematic of acetate metabolism in *E. coli* highlighting the predicted adaptive potential of mutations that might improve acetate metabolism. Green arrows highlight positive regulation, and red lines indicate negative regulation. See also Table S2 and Figures S5, S6, S7, S8.

Five genes that harbored mutations (*rpoD*, *cspC*, *dnaJ*, *ydcI*, and *yheO*) were most likely adaptive because they increase the efficiency of acetate metabolism (Figure 4B). As described earlier, *E. coli* convert acetate to Acetyl-CoA through one of two mechanisms. At high extracellular acetate concentrations (100 mM), cells use the AckA-Pta pathway, while at low concentrations, acetate is scavenged via Acs which has a ∼35-fold higher affinity for acetate uptake (*K_m_* = 7 mM vs *K_m_* = 0.2 mM, respectively)^35^. Our selection environment had relatively low concentrations of acetate (∼17mM), so we can expect metabolism via Acs to be dominant. Indeed, mutations in three genes are expected to increase acetate utilization via Acs, but through different mechanisms. Mutations arose in *rpoD,* which encodes *σ*^70^, the sigma factor responsible for *acs* transcription^44^. Additionally, we identified missense mutations in *cspC*, which may increase *acs* expression because CspC upregulates *rpoS*, which subsequently inhibits the transcription of *acs*^45^, and it has been shown that acetate metabolism is activated in the absence of *rpoS*^46^. Supporting this, Seong *et al*. (2020) found a loss-of-function mutation in *cspC* in response to selection for increased growth in acetate in *E. coli* lacking four genes involved in Acetyl-CoA consumption. Finally, the mutation *dnaJ* H33Y specifically fails to stimulate the ATPase activity of DnaK^47^, and DnaK is strongly linked to acetate metabolism. The absence of DnaK increases acetate consumption at low concentrations by slowing down utilization via the AckA-Pta pathway and facilitating it via Acs^48^, which should be adaptive in our environment. Not related to Acs expression directly, mutations arose in the transcription factor *ydcI* which has been shown to control the carbon flux into the TCA cycle by repressing *gltA*^49^, which is the rate-limiting step when cells are grown solely on acetate^50,51^. Consistent with this, a *ydcI* deletion mutant has been shown to grow faster than WT *E. coli* in acetate^52^. Finally, YheO has been experimentally validated to be a transcription factor^52^, but there is no predicted biochemical characterization or biological function. However, the binding profile of YheO shows that YheO targets the acetate kinase *ackA*^52^, which catalyzes the formation of acetyl phosphate from acetate to ATP. Thus, YheO may activate or repress *ackA*, and mutations in this gene could lead to increased acetate metabolism depending on the function, although this is still unknown.

During growth on glucose, acetate accumulation can be toxic to cells and can inhibit microbial growth which is hypothesized to occur through different mechanisms^28,30,36^. The first explanation is the uncoupling effect of acetic acid (Hac) on the proton motive force, causing excess protons to be pumped out of the cell to maintain membrane potential diverting energy away from growth^53^. Secondly, the presence of acetate anions in the cell increases the internal osmotic pressure^54,55^. For example, 8mM acetate was shown to decrease the growth of *E. coli* by 50% when grown in glucose^55^. In agreement with previous work, mutations arose in our experiment that may contribute to increased acetate tolerance. In addition to acetate metabolism, *ydcI* is important for maintaining pH homeostasis and has been shown to be responsible for acid stress resistance in *Salmonella enterica* and *E. coli*^52^. We also saw the same ∼15-kb deletion in many clones, which leads to the excision of the lambdoid prophage e14 (Figure S6). This e14^−^/*icd^+^* mutation has previously been observed by Moreau and Loiseau (2016) in *E. coli* in response to phosphate starvation conditions, which leads to a build-up of acetic acid in the medium. Excision of the prophage e14 and restoration of *icd^+^* increased resistance to acetic acid stress^56^.

Performing our experiment with six distinct biological replicates (Figure S1) allowed us to examine the degree of repeatability, if any, of phenotypic and genetic adaption. As observed in other “parallel replay experiments” where identical replicate populations evolve under identical conditions^57^, there were many examples of parallelism at the genetic level, where mutations in the same gene, or even at the nucleotide level, independently evolved in multiple chemostat populations, suggesting a strong adaptive benefit (Figure 4). Identical mutations were observed for both *rpoD* (G1339A; chemostats C, D, and E) and *cspC* (A60T; chemostats B and C). Furthermore, the same e14^-^/*icd*^+^ deletion arose independently in four chemostats (A, B, D, and F), which was due to a site-specific recombination in 11-bp repeats in *icdA* and *icdC*, identical to the mutation found by Moreau and Loiseau (2016) (Figure S6). Additionally, beneficial mutations accumulated in many of the same genes (Figure S7). Six unique mutations independently arose in the transcription factor *yheO* across three chemostat populations. Five of these six mutations occurred in the DauR-like helix-turn-helix domain that is likely to act as the DNA-binding domain, potentially affecting transcription of target genes, one of which is *ackA*^52^. Seven independent mutations arose in another transcription factor, *ydcI*, in four different chemostats across six positions, demonstrating additional parallelism at the nucleotide level. Non-synonymous mutations made up all the mutations across the length of the gene (Figure S7). Finally, nine mutations arose across seven sites in three chemostats in *proQ,* a global small noncoding RNA-binding protein that binds to ∼450 RNA targets^58–60^. In comparison to *yheO* and *ydcI* however, mutations in *proQ* were mostly frameshifts and nonsense mutations causing complete loss of the C-terminal domain (Figure S7).

In addition to parallelism, we also observed heterogeneity in the evolutionary response across replicate chemostat populations. Broadly speaking, divergent mutational patterns can be grouped in the following way: 1. chemostat A, 2. chemostats B and C, and 3. chemostats D, E and F (Figure S8). 1. Chemostat A had a unique mutational pattern relative to other chemostats that had signatures of sweeping events, with all clones harboring the same base set of mutations (*ydcI* I226T, *rseA* A156S, and *pyrE* −41 ΔG) and some clones having an additional 1-2 mutations (Figure 4A). 2. Chemostats B and C displayed similar mutational patterns, with all 20 clones having the same mutation in *cspC*, suggesting this was an early mutation that rapidly swept to fixation in both populations. Similar mutations arose in the *cspC* background as the experiment progressed, for example, in *yheO* (Figure 4A). 3. Finally, chemostats D, E, and F showed similar mutational patterns, with many clones harboring mutations in *rpoD*, *proQ*, and *ydcI*. One difference, however, was the *dnaJ* H33Y lineage in chemostat D as this was the only chemostat this mutation arose. Mutational differences across chemostat populations also highlighted signs of epistasis (interactions between mutations) because some mutations appear to occur more often together than others. For example, 5/6 *yheO* variants arose in a strain that had a *cspC* I20M mutation. Furthermore, every *proQ* mutation (nine mutations across three chemostats) co-occurred within a *rpoD* A477T background. Further experiments are needed to validate and evaluate the nature of these epistatic interactions and how they contribute to increased acetate metabolism.

### Genotype-to-phenotype characterization of trade-offs between acetate and glucose metabolism

To identify metabolic trade-offs, we compared clone growth data (relative µ_max_ of clones in acetate and glucose) to genotypes (Figure 5, raw growth data shown in Figure S9). Specifically, we tested if specific mutations or sets of mutations that increase acetate assimilation decrease growth in glucose. There were two clear examples where this was the case. Clones with a *cspC* I20M mutation (all 40 clones from both chemostats B and C) often displayed considerable increases in growth rate in acetate but were accompanied by a significant trade-off in glucose. Specifically, 13/20 clones from chemostat B and 19/20 clones from chemostat C showed significantly reduced growth in glucose relative to the WT (B: one-way ANOVA, F_20,67_=12.2, *p*<0.0001; C: one-way ANOVA, F_19,64_=11.47, *p*<0.0001, pairwise differences assessed with Tukey HSD; Figure 5, see Table S1 for all pairwise comparison statistics). Additionally, clones with the mutation *rpoD* A447T (chemostats D, E and F - Figure 4) often showed decreased growth in glucose (15/31 clones) but unlike *<u>cspC</u>* I20M mutants, didn’t show a significant increase in growth rate in acetate (2/31 clones, Figure 5, Figure S9, Table S1). Other mutations or sets of mutations did not display any measurable metabolic trade-offs in glucose. These included clones with mutations in *ydcI* (chemostats D, E and F), *dnaJ* H33Y (chemostat D), and most clones isolated from chemostat A (Figure 5, see Table S1).

**Figure 5.**
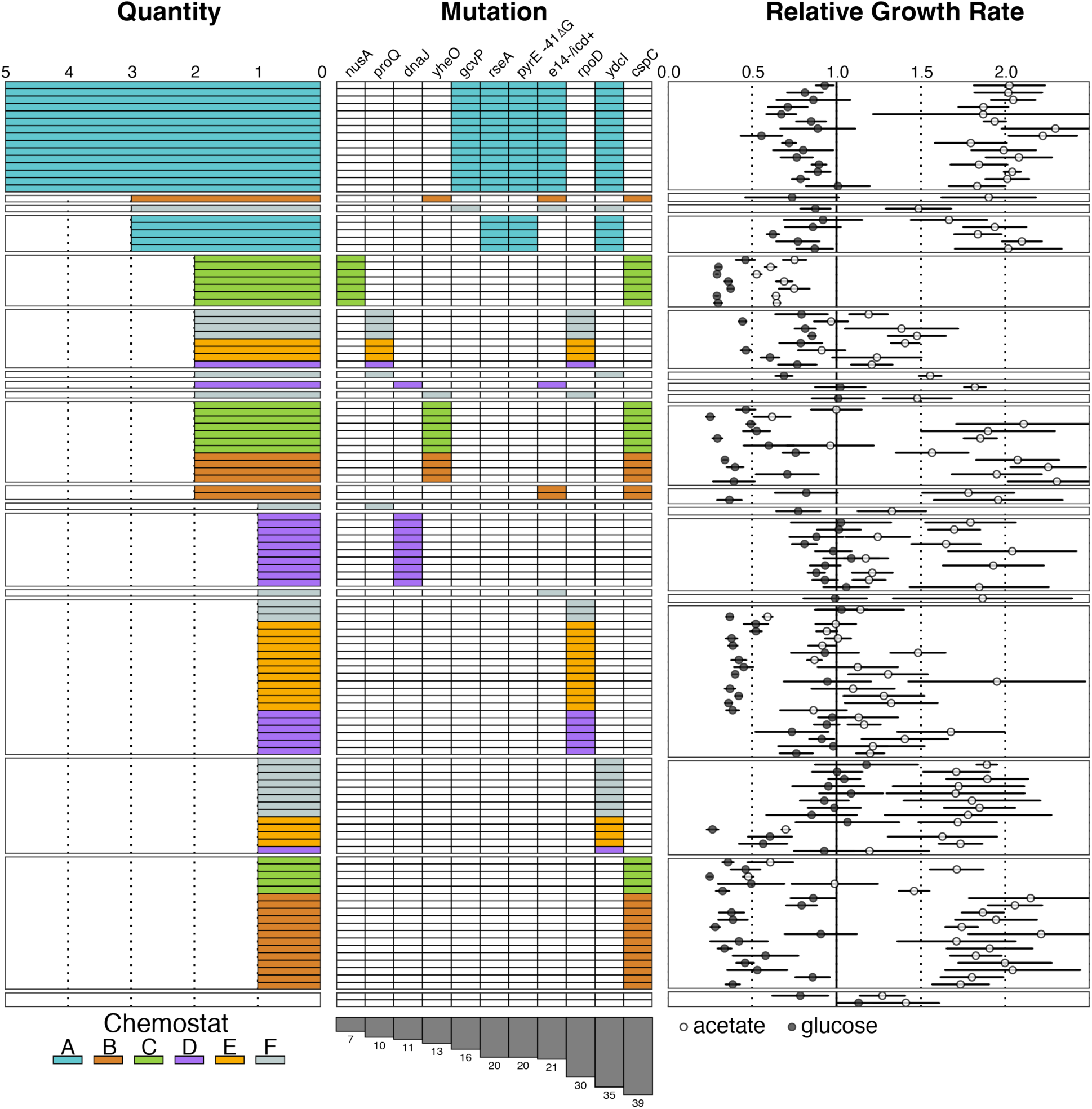
Genotype-to-phenotype map of increased acetate metabolism and its effect on trade-offs in glucose metabolism. We performed an observational analysis to correlate genotype with µ_max_ in acetate and glucose. We organized our data by genotype and the number of mutations, designating chemostat of origin by color. We only included mutations in genes that were found in more than one clone. Mutations in two genes seemed to correlate with a trade-off in glucose medium: *cspC* I20M and *rpoD* A447T. Other genotypes did now show a measurable trade-off in glucose (*e.g*., *dnaJ* H33Y, mutations in *ydcI*, and clones from chemostat A that have 4-5 variants). See also Figure S9, Figure S10, and Table S1.

As described earlier, many evolved clones from chemostat C unexpectedly grew slower than the ancestor in acetate (Figure 3), which was surprising as the whole population grew markedly faster than the ancestor (Figure 2B). Interestingly, most of these clones exhibited the same genotype: *cspC* I20M/*nusA* R258C. To further investigate this, we measured the relative fitness of a single mutant clone (C5 - *cspC* I20M) and a double mutant clone (C6 - *cspC* I20M/*nusA* R258C) isolated from chemostat C against the ancestor using standard competition experiments (direct competition over three days starting at a 1:1 ratio tracking abundance with genomic sequencing - also see Methods). Expectantly, the *cspC* I20M clone that grows faster than the ancestor in monoculture (Figure 6A) also outcompeted the ancestor in direct competition (Figure 6B). However, the *cspC* I20M/*nusA* R258C clone that grows significantly slower than the ancestor in monoculture (Figure 6A, Table S2, *p*=0.008) also outcompeted the ancestor and had a higher fitness than the single mutant clone (Figure 6B). Unexpectedly, when measuring the fitness of the *cspC* I20M/*nusA* R258C clone relative to the *cspC* I20M, the double mutant had a higher fitness than the single mutant (Figure 6B), which could explain why this genotype arose and increased in frequency during the experiment (37% of the clones isolated from chemostat C).

**Figure 6.**
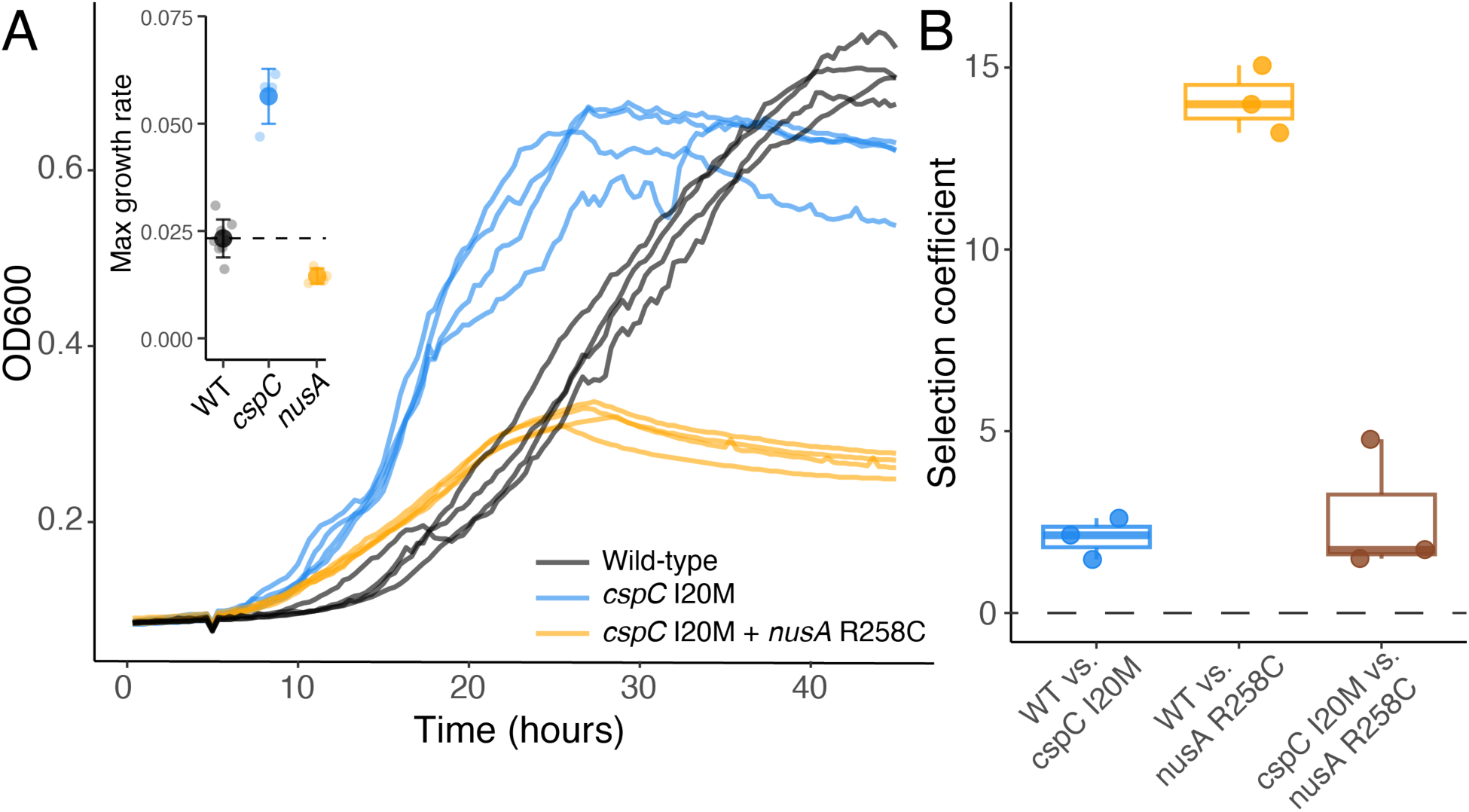
The double mutant *cspC* I20M *nusA* R258C shows reduced growth in acetate but is still competitive against the wild-type and the single mutant progenitor. A) Raw growth curves measured with a 96-well plate reader. Maximum growth rate shown in inset. The *cspC* I20M clone (strain C5) grew significantly faster than the ancestor but the *cspC* I20M/*nusA* R258C clone (strain C6) grew significantly slower with drastically reduced yields. B) The selection coefficient was measured as logarithmic ratio over time of the second listed mutant relative to the first listed mutant in the x-axis (*e.g.*, cspC I20M relative to WT). Thus, a selection coefficient (s) = 0 is equal fitness and positive is increased fitness. Despite significantly reduced monoculture growth in acetate, the *cspC* I20M/*nusA* R258C clone outcompeted both the ancestor and the *cspC* I20M clone.

## Discussion

Evolutionary trade-offs are widespread, and experimental evolution studies have been a powerful approach to directly measure trade-offs and determine the underlying genetic causes^3,8–15^. Adaptive laboratory evolution (ALE) is increasingly being used in industrial settings to select for fast-growing strains, particularly for increased utilization of alternative carbon sources. However, adaptation to one environment often leads to antagonistic pleiotropy for another and understanding the genetics underlying these trade-offs is critical for adaptation of bacteria for bioremediation and biomanufacturing purposes. Here, we utilize continuous cultivation to evolve *E. coli* in media containing acetate as the sole carbon source in turbidostats^39^ (Figure 1C). Acetate is a promising feedstock for industrial applications as it is inexpensive and can be obtained from a number of renewable sources^21,61^. In *E. coli*, however, growth on acetate is slow when compared to glycolytic nutrients such as glucose as it is less preferred with respect to NADH and ATP generation^20^. Having been successful in other studies, ALE is a promising approach to increase bacterial growth and/or yield on acetate^32,37,38^, but trade-offs have not been directly studied in response to increased acetate assimilation. After ∼100 generations of turbidostat evolution, we isolated 119 clones from six independent chemostat populations and measured growth in acetate and glucose separately to determine the response to selection and potential trade-offs. We then characterized the genetic basis for increased acetate assimilation and characterized the genotype-to-phenotype map of acetate adaptation and metabolic trade-offs in glucose.

As expected, all six *E. coli* populations increased in fitness (measured here as maximum growth rate) in our ALE environment: Chi.Bio chemostat reactors with acetate as the sole carbon source (Figure 2B). We then selected 20 random clones from each population to analyze further (19 clones from chemostat C due to the inability to revive the −80°C stock for one clone) and measured the absolute fitness of each clone by calculating the maximum growth rate from growth curves. Like the population-level data, many clones exhibited increased exponential growth relative to the ancestor in acetate (Figure 3, Figure 5). We also characterized trade-offs in glucose by measuring the absolute fitness of all 119 clones in medium containing glucose as the sole carbon source. There was evidence of a metabolic trade-off associated with increased acetate assimilation, with 45% of clones showing significantly reduced growth in glucose (Figure 3, Figure 5, Table S1). 55% of clones, however, showed no measurable trade-off. This is consistent with other experimental examinations of evolutionary trade-offs, where trade-off patterns were often heterogenous amongst measured clones^3,8,62^. For example, in response to high temperature (42.2 °C) adaptation in *E. coli*, trade-offs were common but variable at lower temperatures, with 31% and 56% of clones exhibiting trade-offs at 37°C and 20°C, respectively^3^.

Fitness trade-offs can arise from antagonistic pleiotropy where beneficial mutations in the evolution environment are maladaptive in another, mutation accumulation where neutral mutations can arise in the selective environment that decrease fitness in an alternative environment, or some combination of both. Our ALE experiment was short at roughly 100 generations in length, so we expected that antagonistic pleiotropic effects of early adaptive mutations were more likely to contribute to trade-offs in glucose. Indeed, this is what we saw for some adaptive mutations. For example, *cspC* I20M alone resulted in an 85% increase in growth rate in acetate (Figure 5, Figure S7) and 17/25 of the fastest-growing clones had this mutation (Table S1). Furthermore, in direct competition with the ancestor, a *cspC* I20M mutant had a selection coefficient of 2.1 (Figure 6), suggesting that this mutation had a large adaptive benefit. In line with this, we found this mutation in all 20 clones isolated from chemostats B and C (Figure 4), indicating that it arose and quickly swept to fixation. A mutation in *cspC* is predicted to be adaptive in acetate because CspC positively regulates *rpoS* and increases its stability^45^, and acetate metabolism is activated in the absence of *rpoS*^46^. However, a *cspC* I20M mutation also resulted in a large trade-off for growth in glucose: *cspC* I20M clones grew 57% slower than the WT and comprised 12/25 of the slowest-growing clones, suggesting that this mutation generates trade-offs via antagonistic pleiotropy.

On the other hand, other mutations that had a large adaptive benefit in acetate displayed no measurable trade-off in glucose. Clones harboring *dnaJ* H33Y isolated from chemosotat D (Figure 4) grew twice as fast as the WT in acetate and 7/25 of the fastest-growing clones had this genotype. It has been shown that a mutation at this amino acid residue results in the failure of DnaJ to stimulate DnaK ATPase activity^47^, and in the absence of DnaK activity, acetate metabolism is switched to the Acs pathway which allows for faster growth at lower concentrations of acetate, which should be adaptive in our environment^48^ (17.3 mM acetate). Unlike *cspC* I20M and consistent with Angles et al. 2017, clones with mutations in *dnaJ* did not show reduced growth in glucose (Table S1). These results are consistent with other experiments and models that suggest that the fitness effects of adaptive mutations in the selected environment can have a range of pleiotropic effects in an alternative environment^3,8,11^. However, our results suggest that some early adaptive mutations (<100 generations) can have significant pleiotropic effects in non-selected environments, which is inconsistent with the observation that antagonistic effects of adaptive mutations occur more often over longer evolutionary timescales^11,13^. It is possible, however, that evolution over longer timescales in this system will result in further acetate specialization that could be accompanied by further decreases in fitness for growth in other carbon sources.

Surprisingly, a handful of clones from chemostat C grew significantly slower than the WT, which was unexpected as the turbidostat leads to a steady-state environment in which nutrients are in abundant supply and the growth rate of cells is the limiting factor, creating a strong selective pressure for faster-growing mutants^41,42^. Whole-genome sequencing revealed that many of these clones had the same genotype (*cspC* I20M/*nusA* R258C; Figure 5), suggesting they were isolated from the same lineage. To investigate this genotype further, we examined the relative fitness of a double mutant (strain C5 – *cspC* I20M/*nusA* R258C) against the WT and the presumed single mutant progenitor (strain C6 – *cspC* I20M). The double mutant had a higher fitness than both the WT and the single-mutant clone, explaining its ability to increase in frequency once it arose (7/19 clones from chemostat C). This could be due to a cross-feeding interaction that arose in the *nusA* R248C genotype, allowing it to grow on the metabolic byproducts of the *cspC* I20M strain. Cross-feeding interactions have evolved in other experimental evolution studies founded by a single clone that led to a stable coexistence between two or more genotypes^63,64^. However, this finding highlights how monoculture growth rates can be a poor predictor of population fitness^65^. Nevertheless, absolute fitness can be assessed in a high-throughput manner and calculating growth parameters from growth curves, such as growth rate, yield, and the area under the curve (AUC; Figure S10), can be utilized for evaluating large sets of phenotypes, especially in industrial settings where faster growth is the desired result^66^, as in our ALE experiment here.

Identifying repeated targets of selection, especially at the gene or pathway level, is common during experimental evolution across replicate microbial populations^67–69^. We also observed parallelism across chemostat populations during our ALE (Figure 5, Figure S6, Figure S7), suggesting strong selection for those genes for increased growth and/or tolerance in acetate. Of note, we observed parallelism at the nucleotide-level in *cspC* A60T and *rpoD* G1339A, with these mutations independently arising in two and three different chemostats, respectively. CspC and RpoD are linked to acetate metabolism as they can influence Acs expression^37,44–46^ (Figure 4B) explaining an adaptive benefit for mutations in these genes. Repeated mutations were also observed at the gene level, with non-synonymous mutations arising in three genes, *ydcI*, *yheO*, and *proQ,* in multiple chemostats (Figure S7). The transcription factor YdcI has been shown to be important for acetate metabolism because it represses *gltA*^52^, which when induced is the rate limiting step for the TCA cycle when grown in acetate^50,51^ (Figure 4B). However, mutations in *yheO* and *proQ* have not been reported in *E. coli* evolved in acetate. YheO is a transcription factor for which the biological function is unknown. In relation to acetate metabolism, though, Gao et al. 2018 showed the YheO binds to *ackA* in glucose medium. If we map the non-synonymous mutations onto *ydcI*, mutations arose in the dauR-like HTH transcriptional binding domain (Figure S7), suggesting that mutations in this region may affect regulation of *ackA,* but the mechanism is unclear. ProQ is a global small non-coding RNA-binding protein that associates with ∼450 transcripts involved in a diversity of biological functions^59,60^. Mutations in this gene were presumably loss-of-function mutations including nonsense and indels leading to complete or partial loss of the linker as well as complete loss the C-terminal domain (Figure S7). Despite the lack of a clear explanation, the observation of repeated mutations in *yheO* and *proQ* suggest a strong adaptive benefit in *E. coli g*rowing solely in acetate, and current work is underway to identify the mechanism(s) via transcriptomics.

We also observed divergent mutational responses to selection across replicate chemostat populations. In laboratory evolution experiments, it is not unusual for replicates with similar phenotypic characteristics to have mutational differences, and these differences can contribute to phenotypic and fitness differences in other environments^70,71^. Here, chemostat populations displayed different mutational patterns and grouping our data by genotype (Figure 5) highlights these. For example, many clones isolated from chemostats B and C showed similar genotypes (green and orange grouped together in Figure 5), with many mutations arising in *cspC* and *yheO*. Clones isolated from chemostats D, E, and F displayed similar genotypes (yellow, purple, and grey grouped together in Figure 5), with mutations commonly occurring in *ydcI*, *rpoD*, and *proQ* (Figure 4A). This also highlighted the co-occurrence of different mutations. For example, mutations in *yheO* co-occurred with *cspC* I20M and mutations in *proQ* always co-occurred with *rpoD* A447T (Figure 4A, Figure 5). This could be due to epistatic interactions between these gene pairs, which could be important as interactions among sets of mutations can alter the mutational trajectory of an evolving population. Altogether, these subtle differences suggest that there could be multiple adaptive routes to increased acetate metabolism in *E. coli* and these divergent paths could result in diverse impacts on fitness in glucose or environments with other carbon sources.

Acetate has gained much attention as a promising low-cost feedstock to develop economically feasible bioprocesses for obtaining desired bioproducts^20,21,61^. *E. coli* has been extensively explored for this purpose as it can use acetate as a sole carbon source. However, growth of *E.coli* on acetate is slow relative to other preferred carbon sources such as glucose, so ALE has the potential to be a useful tool to produce improved phenotypes for acetate utilization. Acetate, however, is abundant in mixed feed systems that contain many different carbon sources, such as lignocellulosic hydrolysates, a source of biomass to produce a range of bio-products^72^. Therefore, co-utilization of acetate and other carbon sources is important for efficient bioproduction^73^. Yet, we have shown that ALE for faster growth and tolerance in acetate can quickly result in trade-offs for glucose utilization, the most abundant hexose in lignocellulosic hydrolysates^74^. This could affect the potential for co-utilization of different carbon sources in complex feedstocks that contain acetate, reducing the efficiency of the system. Overall, our results suggest that directed evolution within homogeneous environments for alternative carbon sources can rapidly result in trade-offs for other carbon source metabolism due to the antagonistic pleiotropic effects of adaptive mutations. This may limit the utility of ALE in homogenous environments for strain improvements in biotechnology involving complex or dynamic nutrient conditions.

## Materials and Methods

### Strains and growth conditions

We used *E. coli* BW25113 for all experiments. Bacteria were cultured in minimal media [MM – recipe in Table 1;^75^] with 2 g/L sodium acetate (resulting in an acetate concentration of 17.3 mM) as the sole carbon source for adaptive laboratory evolution (ALE) in chemostats as well as growth rate measurements. Additionally, MM with 25mM glucose was used for growth rate measurements.

**Table 1.** Recipe for minimal media.

| Minimal media | Stock concentration | Volume (per liter) | Final concentration |
| --- | --- | --- | --- |
| Potassium phosphate buffer (see below) | 0.1 M | 100 mL | 10 mM |
| MgSO<sub>4</sub> | 1.5g/50mL | 2 mL | 0.5 mM |
| CaCl<sub>2</sub> | 34 mM | 1 mL | 0.034 mM |
| NH<sub>4</sub>Cl | 8 M | 1 mL | 8 mM |
| Disodium EDTA | 7.44 mM | 1 mL | 7.44 μM |
| FeSO<sub>4</sub>·7H<sub>2</sub>O | 0.10g/50mL | 0.5 mL | 3.6 μM |
| Trace elements (see below) |  | 5 mL | N/A |
| Trace elements | Stock concentration | Volume (per liter) | Final concentration |
| ZnSO<sub>4</sub>·7H<sub>2</sub>O | 0.5 g/50mL | 1 mL | 0.17 μM |
| MnCl<sub>2</sub>·4H<sub>2</sub>O | 0.15 g/50mL | 1 mL | 0.08 μM |
| H<sub>3</sub>BO<sub>3</sub> | 1.5g/50mL | 1 mL | 2.43 μM |
| CoCl<sub>2</sub>·6H<sub>2</sub>O | 1g/50mL | 1 mL | 0.42 μM |
| CuCl<sub>2</sub>·2H<sub>2</sub>O | 0.1g/50mL | 0.5 mL | 0.03 μM |
| NiCl<sub>2</sub>·6H<sub>2</sub>O | 0.1g/50mL | 1 mL | 0.04 μM |
| NaMoO<sub>4</sub>·2H<sub>2</sub>O | 0.15g/50mL | 1 mL | 0.06 μM |
| Potassium phosphate buffer (0.1M) | Amount (per liter) |  |  |
| KH<sub>2</sub>PO<sub>4</sub> | 4.14 g |  |  |
| K<sub>2</sub>HPO<sub>4</sub> | 12.119 g |  |  |

### Adaptive laboratory evolution for increased acetate metabolism in Chi.Bio chemostats

To select for increased acetate metabolism in *E. coli*, we utilized Chi.Bio’s all-in-one automated chemostat culturing platform^39^. Six Chi.Bio reactors were set-up as per the instructions in Steel et al. 2020 and detailed at https://chi.bio/operation/. Additionally, we laser cut stands to elevate the reactors to keep the liquid level in the reactors above the liquid level in the fresh media and waste containers (this prevents liquid draining into the reactor itself in the event a pump makes a poor seal). The stand also framed the reactors to keep them stationary during the experiment (Figure 1A). To initiate the chemostat cultures, *E. coli* BW25113 was grown overnight in 50mL of MM + 2g/L sodium acetate from a single colony on an agar plate, shaking at 200rpm at 37°C. Next, we made a 1:100 dilution of the overnight culture in 20mL of fresh media in six test tubes (30-mL Fisherbrand™ Class A Clear Glass Threaded Vials with Caps Cat #14-955-320) that were then inserted into a reactor that had been previously blanked with a test tube of fresh media, setting the OD zero point ^39^. The experiment was run through the Chi.Bio web interface with the following parameters: turbidostat mode (OD maintained at approximately 2% of its set point) with an OD set point of 1.0, 37°C, and a stir rate of 0.5 (Figure 1B). It ran for about 12 days, corresponding to roughly 90 generations of *E. coli* evolution (Figure S1). 10% glycerol cryogenic stocks were made of the final timepoint for each chemostat and stored at −80°C.

### Growth rate of evolved populations in in Chi.Bio chemostats

We measured the growth rate of the wild-type ancestor and the six chemostat evolved populations (A-F) using the Chi.Bio chemostats with the OD set to follow a dithered waveform that rapidly dilutes the culture to a lower OD once the user-defined upper limit is reached, allowing for a precise measurement of growth rate ^39^. Specifically, we grew the wild-type and evolved chemostat populations from cryogenic stocks overnight in 15 mL culture tubes (Cat# 352057) with 1mL of MM + 2g/L sodium acetate, shaking at 200rpm at 37°C. Cultures then were inserted into the chemostats were prepared as described above. The experiment ran with an OD set-point of 1.0 set to a dithered waveform (OD 1.0 ± 0.04), 37°C, and a stir rate of 0.5 (Figure 2A, Figure S3).

To determine the growth rate, we developed a custom R script called *chemostat_regression*^43^. Using the library argparse, measurements describing the chemostat optical density, fresh media flow rate, and waste media flow rate over time is input as a data frame from a .csv or .tsv file. Additionally, tunable parameters for adjusting calculations (minimum cycle timepoint, local range for finding relative optical density maxima and minima) are input, as well. Growth cycle ranges are identified via timepoints with non-zero fresh- and/or waste-media flow rates. Growth rate is calculated across each cycle range by fitting a linear model of optical density in response to time (function = lm(); formula = OD ∼ time). The resulting linear model slope predictions were then visualized via ggplot2 (Villanueva and Chen 2019, Figure 2A). The *chemostat_regression*software is available on github at https://github.com/lanl/chemostat_regression.

### Growth rate measurement of single strains clones

Twenty clones from each evolved population (19 clones from chemostat C) were randomly isolated by diluting overnight cultures in a 1.5mL microcentrifuge tube and plating them at a density of 100-200 colonies per plate (MM + 2g/L sodium acetate + 15g/L agar) using glass beads. Twenty colonies were haphazardly picked, re-plated, then single colonies were picked again to ensure clonality. All clones were grown overnight in 15mL cultures tubes with 1mL MM + 2g/L sodium acetate, and 10% glycerol cryogenic stocks were made and stored at −80°C.

We measured the growth rate of clones using a BioTek Epoch 2 Microplate Spectrophotometer in MM + 2g/L sodium acetate and MM + 25mM glucose to determine if evolved clones exhibited a trade-off between increased acetate metabolism and glucose metabolism (their preferred carbon source). Samples (wild-type ancestor and evolved clones) were prepared by plating glycerol stocks on agar plates with either acetate or glucose depending on the carbon source being measured. Plates were grown for 24h at 37°C after which 1 mL cultures in 15mL culture tubes were initiated from plates and grown overnight in MM + 2 g/L sodium acetate or 25mM glucose at 37°C, shaking at 350rpm. Then, each sample was diluted 1:100 in 1 mL fresh media and 200µL was placed into 4 replicate wells (8 replicates for wild-type) in a Nunc™ Edge™ 96-cell well anti-evaporation plate (Cat# 167574). Plates were run on the Epoch 2 for 48h at 37°C, shaking at 580 cpm (orbital), with the OD600 being measured every 20 minutes. The maximum growth rate (µ_max_) of each sample was calculated using gcplyr (https://github.com/mikeblazanin/gcplyr/), a model-free, non-parametric analysis software for microbial growth curves^77^.

Single-strain clones from each chemostat were measured on separate plates, so to rule out potential plate effects that might affect the growth rate measurements, we first compared the µ_max_ of the wild-type measured between each plate as each plate had eight biological replicates of wild-type as a control which was further used to calculate the relative increase in µ_max_ (Figure 3, Figure S4). There was little to no significant difference in the wild-type measurements between plates in both the acetate and glucose environment (Figure S11A,B; acetate - one-way ANOVA, F_5,42_=3.02, *p*=0.020, pair-wise difference assessed with Tukey HSD; glucose - one-way ANOVA, F_5,42_=1.95, *p*=0.106, pair-wise difference assessed with Tukey HSD), suggesting little to no plate effects. Furthermore, we measured the µ_max_ of a selection of clones from each chemostat that exhibited a range of µ_max_ in acetate in the same plate as described above. This showed little difference in the µ_max_ calculated when measured in separate plates (Figure S11C) vs. the same plate (Figure S11D), further confirming the absence of plate effects with this analysis.

### Sequencing and bioinformatic analysis

We sequenced the genomes of all 119 evolved clones as well as 5 clones isolated from the wild-type to determine if there were any mutations relative to the reference *E. coli* BW25113 genome in our starting populations. Genomic DNA was extracted using a Zymo Quick-DNA Fungal Bacterial Miniprep kit (Cat# D6005). Extracted gDNA was quantified using a Qubit 2.0 using the dsDNA High Sensitivity Kit and gDNA length was determined using an E-gel™ Power Snap Electrophoresis system. Libraries were prepared using the NEBNext® Ultra™II Library Prep Kit. Prior to pooling, samples were quantified by running on an Agilent 4200 TapeStation System. Samples were then pooled, cleaned up with PCR AMPure Beads, and then quantified using a Qubit 2.0 and a Bioanalyzer. Finally, the pooled library was run on a NextSeq 2000 using the NextSeq 2000 P3 300 cycle kit (Cat #20040561). Mean coverage across the genome was 1239X for clones from the ancestor and chemostats A, C, and D, and 680X for chemostats B, E, and F. This was due to splitting samples between two different sequencing runs.

To identify *de novo* mutations in ancestral and evolved genomes, we first filtered the reads with trimgalore (https://github.com/FelixKrueger/TrimGalore). Then, we aligned reads to the *E. coli* BW25113 reference genome using the Burrows-Wheeler Aligner algorithm BWA-MEM with the default parameters^78^. Next, we obtained and manipulated .bam files using the Samtools tools suite^79^. Duplicates reads were marked and removed using samtools markdup. Variants were called using both the Genome Analysis Toolkit’s (GATK) HaplotypeCaller^80^ and DeepVariant^81^. Variants were manually validated by visualization using the Integrated Genomics Viewer^82^. Finally, validated variants were pooled and annotated using SnpEff^83^.

### Competition experiment with *nusA* mutant

To elucidate the competitive dynamics with *nusA* mutants that display reduced growth relative to the wild-type, we performed a direct competition experiment with the wild-type ancestor as well as single *cspC* mutant. To initiate competitions, strains C5 (fast grower – *cspC*), C6 (slow grower – *cspC* + *nusA*), and the wild-type *E. coli* BW25113 were plated onto acetate agar plates (MM 2g/L sodium acetate + 15g/L agar) and grown for 24h at 37°C. Then, overnight cultures were initiated by inoculating a colony into 1mL MM + 2g/L sodium acetate in 15mL culture tubes. Cultures were then diluted to an OD600 of 0.1 to normalize biomass. Competitions were started by mixing equal volumetric ratios of competing strains, then diluting 1:100 into 20mL cultures of MM + 2g/L acetate in 150mL flasks. Every 24 hours for 2 days, a 1:100 dilution was inoculated into a fresh 20mL culture. To measure fitness, we used whole-genome sequencing to determine the ratio of different strains over the course of the competition. Genomic DNA was extracted and sequenced at the beginning (time0) and end (time2) of the competition as described above. We analyzed the WGS data to determine the variant allele frequency of specific mutants using the same bioinformatic workflow as described above. The selection coefficient (s) was calculated as:

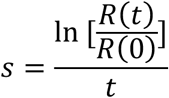

where *R* is the ratio of the mutant to the reference and *t* is the number of generations of the population during the experiment. This means that *s* = 0 is equal fitness, positive is increased fitness, and negative is decreased fitness.

## Acknowledgements

We would like to thank Cheryl Gleasner and Erin Box for performing the whole genome sequencing. This work was supported by the U.S. Department of Energy through the Los Alamos National Laboratory. Los Alamos National Laboratory is operated by Triad National Security, LLC, for the National Nuclear Security Administration of U.S. Department of Energy (Contract No. 89233218CNA000001). This work was funded by a Los Alamos National Lab Director’s Postdoctoral Fellowship to J.T.P. (20230859PRD3). This work has been cleared for public release by Los Alamos National Laboratory (LA-UR-24-32306).

## Author contributions

Conceptualization: J.T.P., B.T.H. and E.R.H.; Methodology: J.T.P. and E.R.H.; Software: S.I.K. and E.A.M.; Formal analysis: J.T.P. and K.T.Y.M.; Investigation: J.T.P.; Writing – Original Draft: J.T.P.; Writing – Review & Editing: All Authors; Visualization: J.T.P.; Supervision: B.T.H. and E.R.H.; Funding acquisition: B.T.H. and E.R.H..

## Supplementary Information

**Figure S1.**
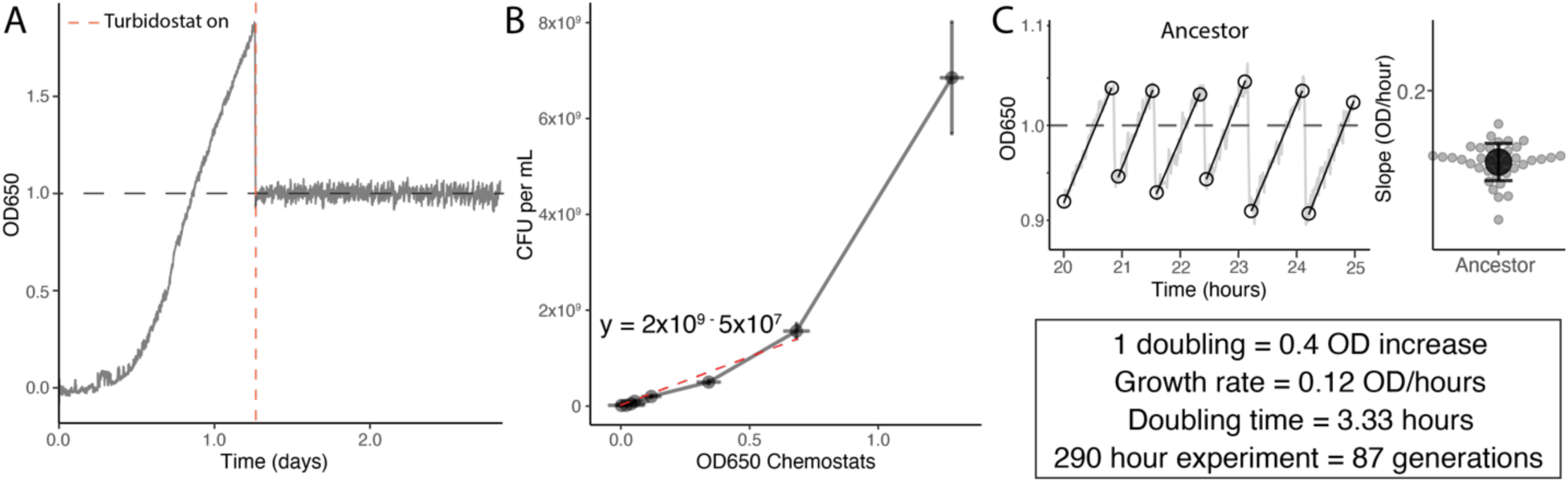
Experimental parameters and calculating number of generations. A) Growth curve of *E. coli* BW25113 in MM + 17.3 mM acetate. We allowed *E. coli* to grow without the addition of fresh media, similar to batch culture, to determine the optical density (OD) near maximal growth rate of cells to inform the turbidostat ALE. We set the OD to 1.0 during turbidostat evolution as this was near the maximal growth rate during exponential growth. B) To determine the number of generations that occurred in the experiment, we first measured colony forming unit (CFU)/mL vs. chemostat OD650 curve to calculate the increase in OD650 the corresponds to one cell doubling, which was 0.4 OD. Like spectrophotometers, the OD650 in chemostats seems to saturate at higher cellular densities, so we used the first 7 datapoints to determine the relationship between CFU/mL and OD650. C) We then used the maximum growth rate measured in chemostats with the OD set to follow a dithered waveform that showed the maximum growth rate of ancestral *E. coli* BW25113 was 0.12 OD/hour. From there, we were able to determine the doubling time of cells in our ALE environment and calculated the number of generations over a 12.5-day (290-hour) experiment to be ∼87 generations. This may be a low estimate as generation time increased during the ALE experiment.

**Figure S2.**
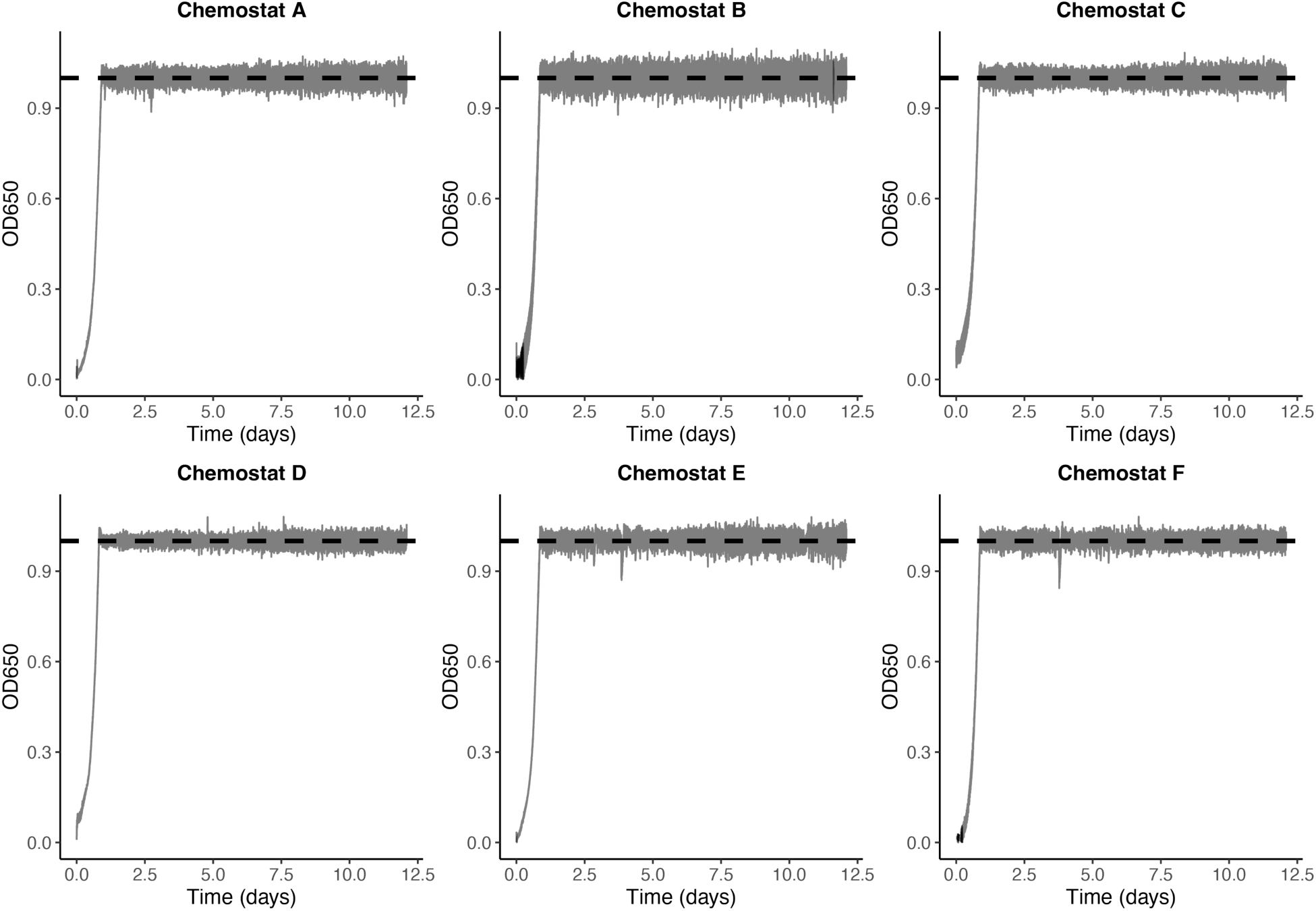
Raw experimental output of all six chemostat from 12.5-day long ALE experiment. Optical density was maintained with 2% of the OD650 set point of 1.0 throughout the whole experiment following initial growth.

**Figure S3.**
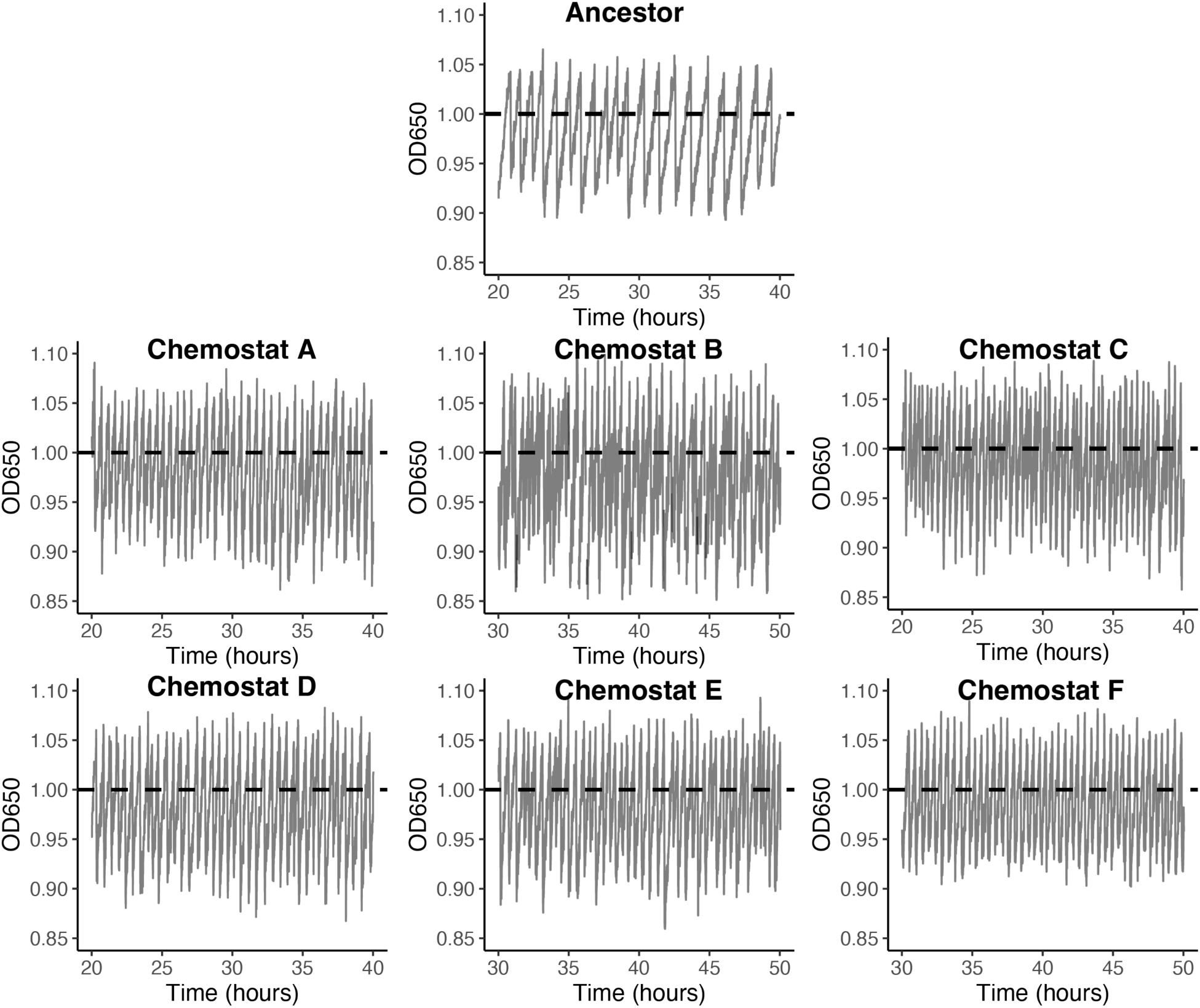
We measured the growth rate of the ancestor and evolved populations in acetate medium using Chi.Bio chemostats with the OD set to follow a dithered waveform. We calculated the slope in between dilutions over a 20-hour window using our custom R script *chemostat_regression*.

**Figure S4.**
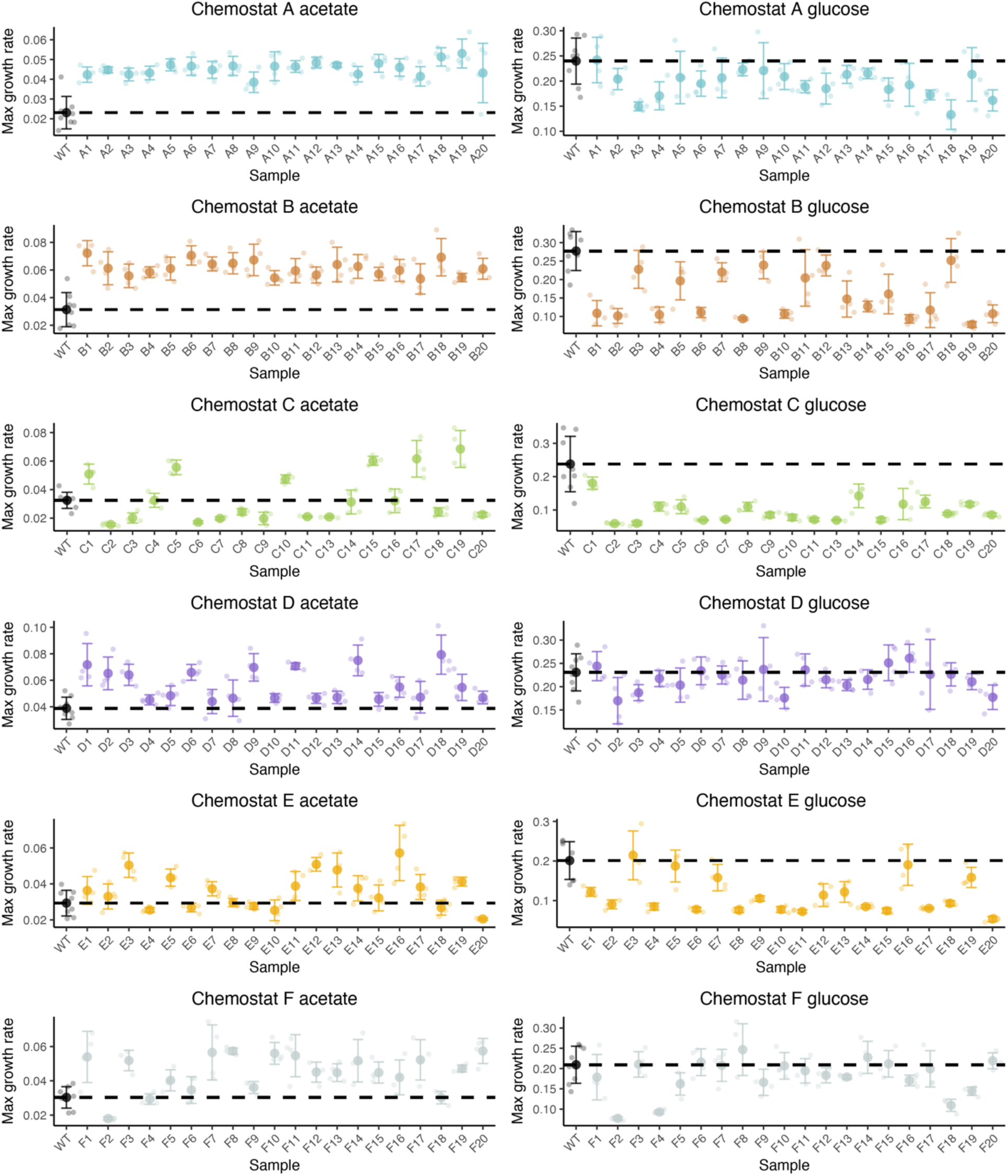
Raw growth rate data from plate reader experiments measuring maximum growth rate (µ_max_) in acetate and glucose media. Calculated maximum growth rate for all evolved clones isolated. The growth of each clone was measured in a plate reader in M9 minimal media containing either acetate or glucose as the sole carbon source. Each clone was measured with four biological replicates. For each measurement, a reference WT *E. coli* BW25113 was measured to calculate fold-change in growth rate (black dotted line) which was used to generate Figure 3.

**Figure S5.**
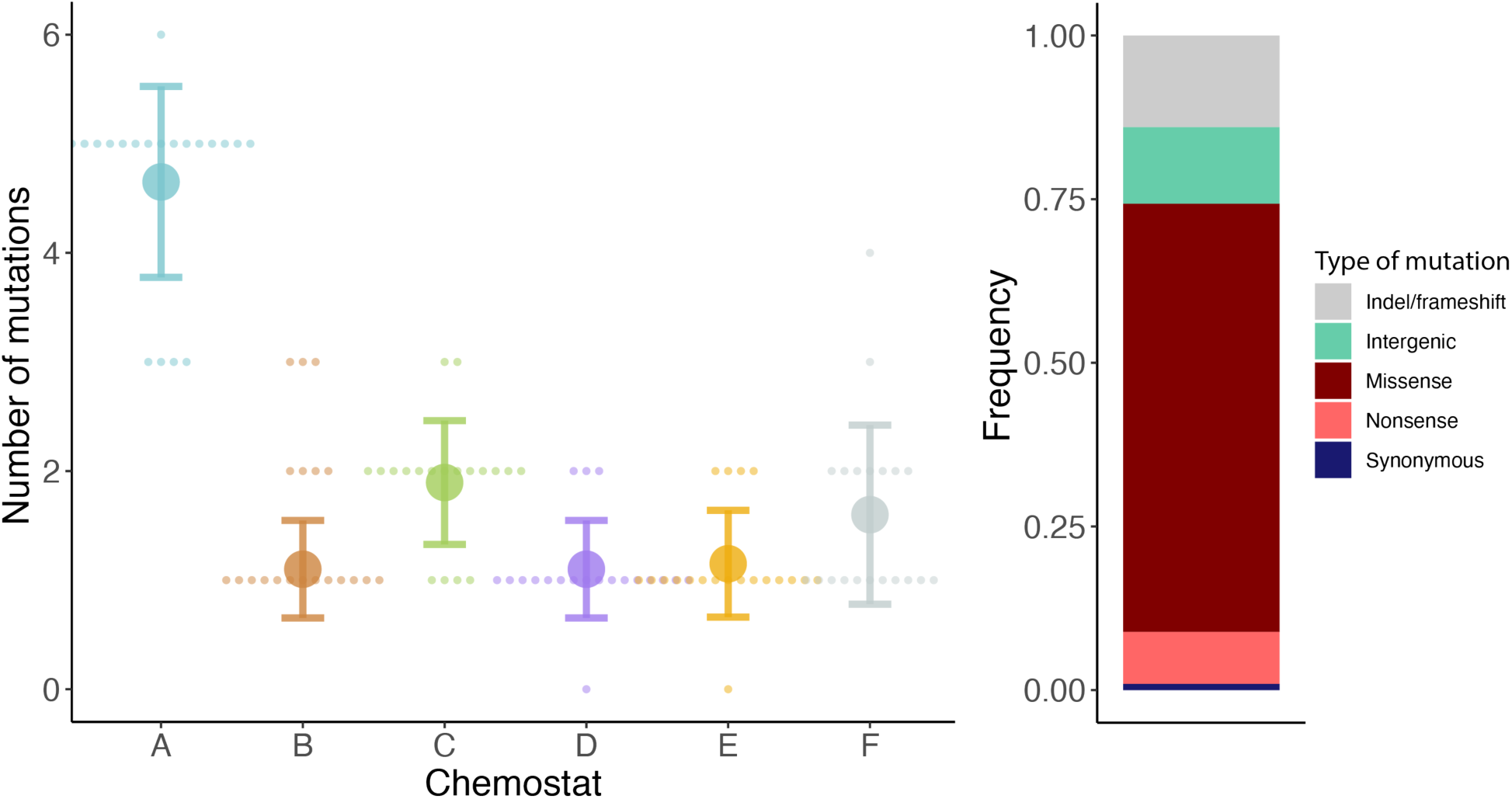
Number and type of mutation that was found after ALE. A) Many clones had only 1-2 mutations, apart from chemostat A, consistent with a short 90 generation experiment. B) Most mutations were non-synonymous mutations, indicating strong selection for faster growth in acetate media.

**Figure S6.**
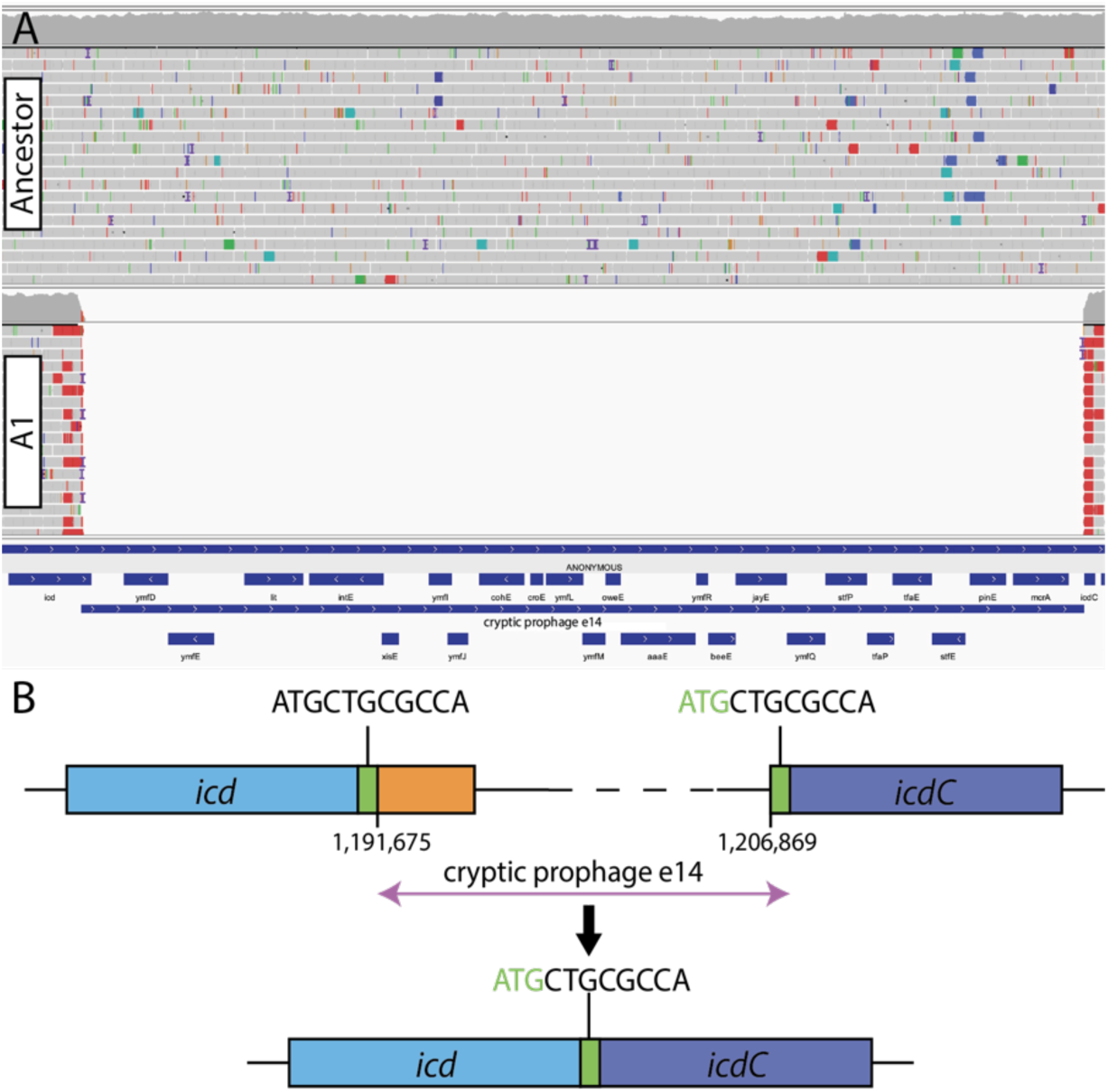
e14^-^/*icd^+^* deletion independently evolved in three chemostats. A) e14-/*icd^+^* mutation consisted of a large 15,193-bp deletion that excised the cryptic prophage e14. Shown is alignments of the ancestor BW25113 and evolved clone 1 from chemostat A. B) Excision occurred through a site-specific recombination of an 11-bp repeat that flanked the prophage (shown in green).

**Figure S7.**
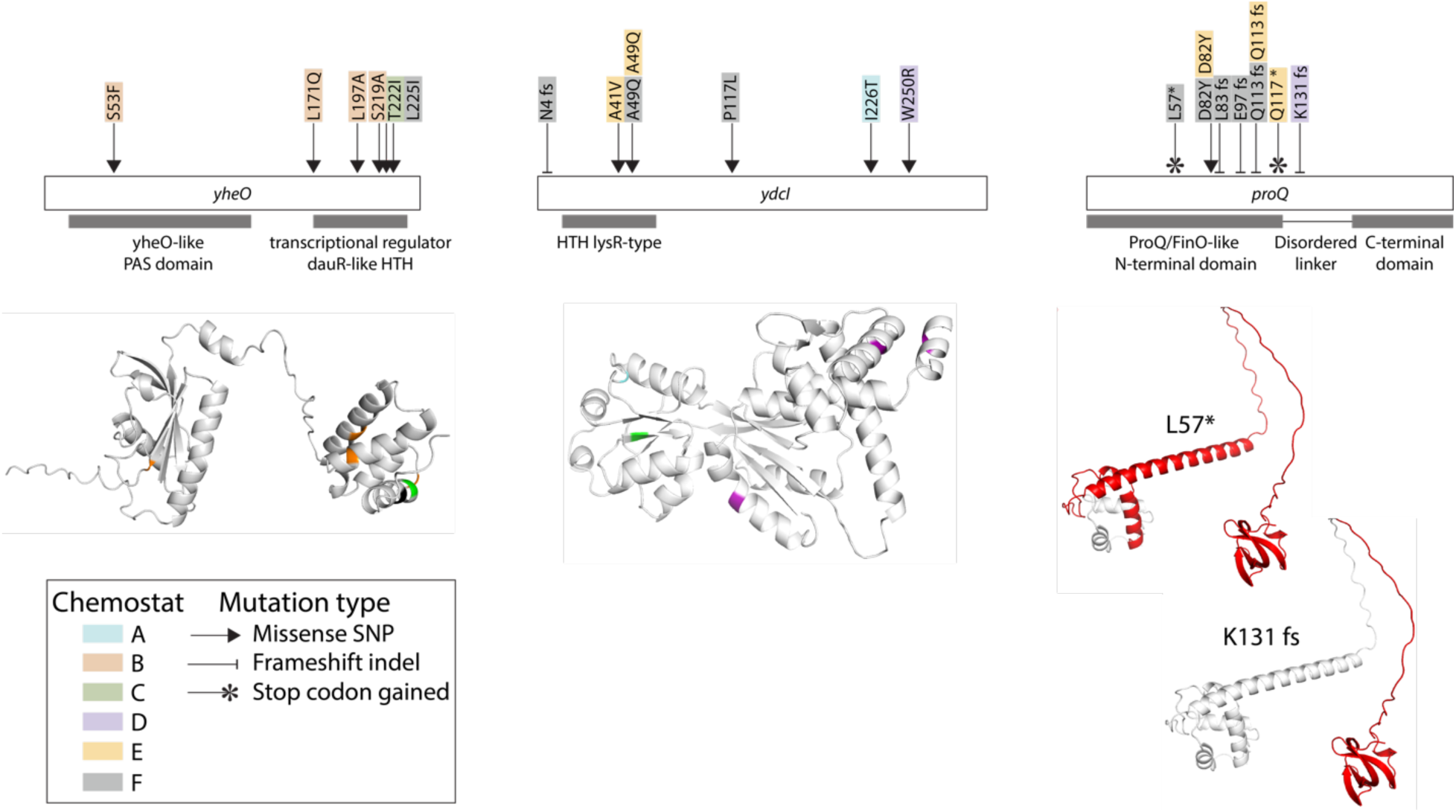
Genes that had a high degree of parallelism. DNA-binding transcriptional regulators YheO, YdcI, and small noncoding RNA-binding protein ProQ all displayed a high degree of parallelism in response evolution in medium containing acetate as the sole carbon source. Mutations and type of mutation are denoted by arrow type, and the chemostat of origin are shown above genes. Protein structures are predicted structures from Alpha Fold and mutations are shown in the proteins. For *proQ*, the earliest and latest mutations are shown that delete part of the gene, displaying deletion of the C-terminal domain (shown in red on the protein structure).

**Figure S8.**
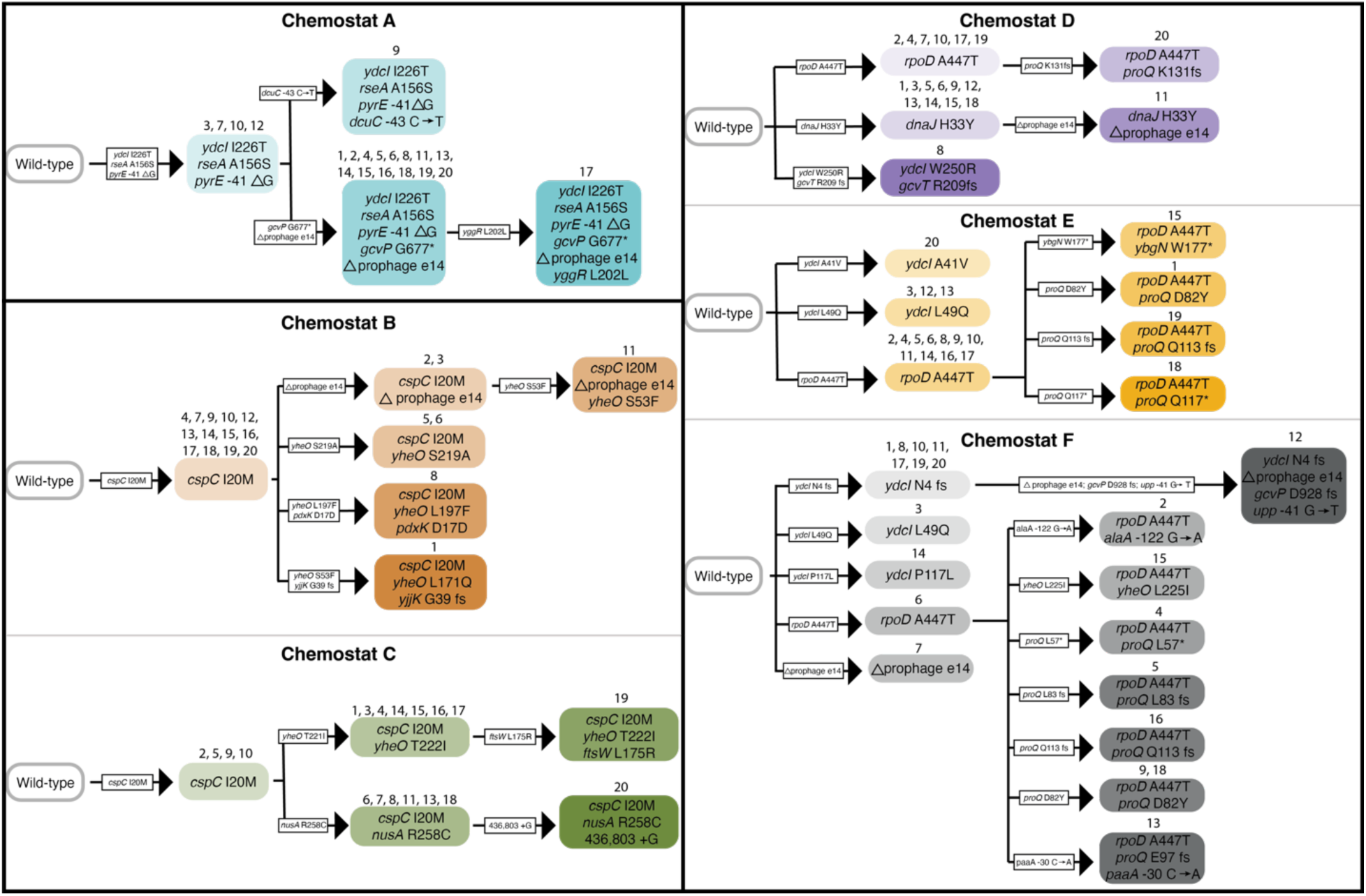
Predicted mutational trajectories of chemostat populations. Based on the presence or absence of mutations, we predicted the mutational trajectories for each chemostat. Response to selection at the genetic level was heterogeneous, with different chemostats displaying divergent mutational patterns. Divergent patterns are grouped by darker boxes (1. chemostat A, 2. chemostats B and C, and 3 chemostats D, E, and F). Divergent mutational patterns also highlight potential epistatic interactions between genes as some mutations commonly co-occur (e.g., *rpoD* and *proQ*; *cspC* and *yheO*). Furthermore, mutational pathways show signatures of sweep events (e.g., *cspC* I20M mutation in chemostats B and C).

**Figure S9.**
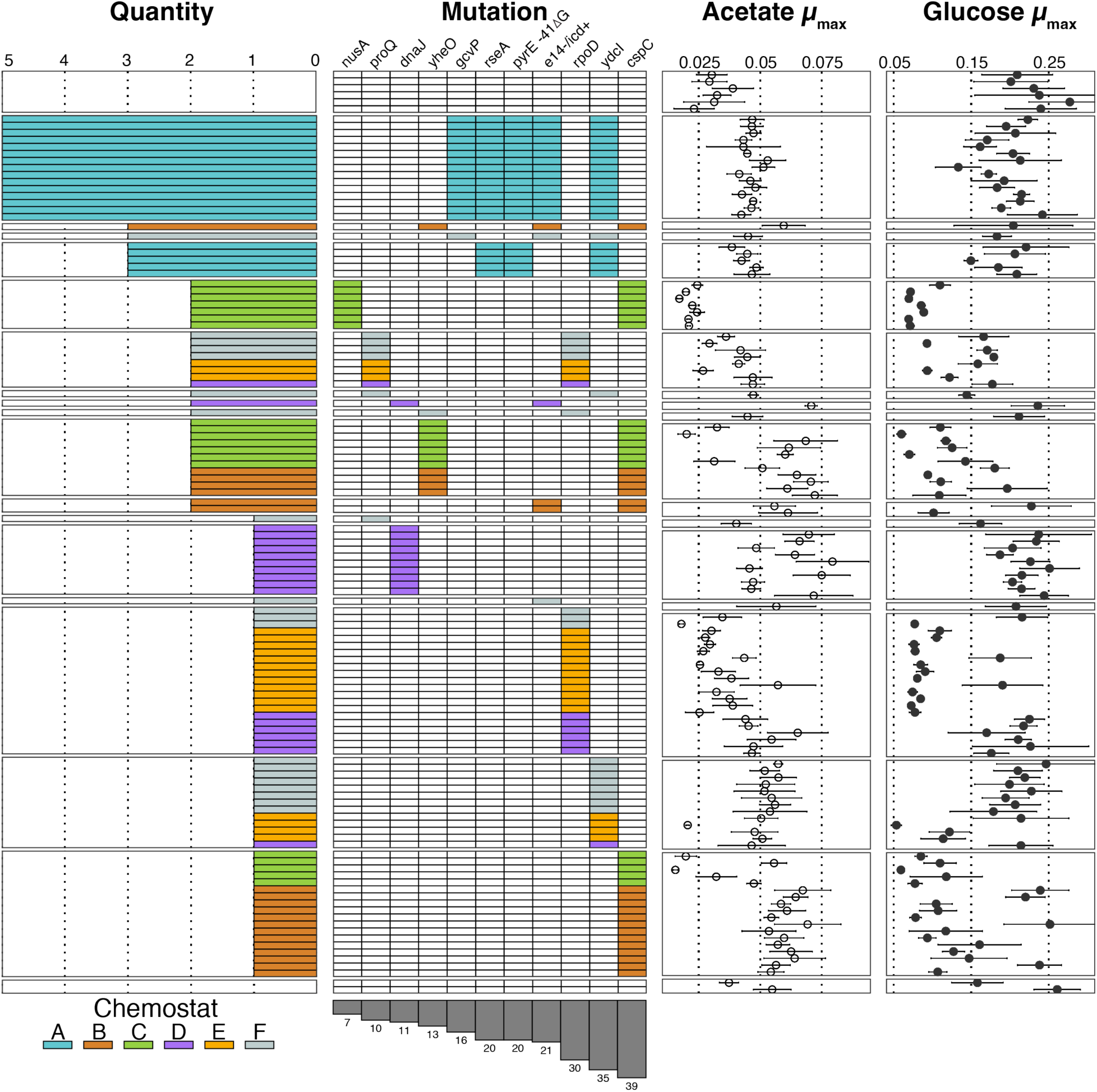
Raw data from growth curves and the genotype-to-phenotype map. Genotype-to-phenotype map of mutations with raw growth rate data compared to relative data shown in Figure 5. For acetate metabolism, large increases in acetate assimilation occurred in clones with mutations in *cspC* and *dnaJ*, suggesting mutations in these genes have a large effect. Wild-type values from each measured plate plotted in the top panel.

**Figure S10.**
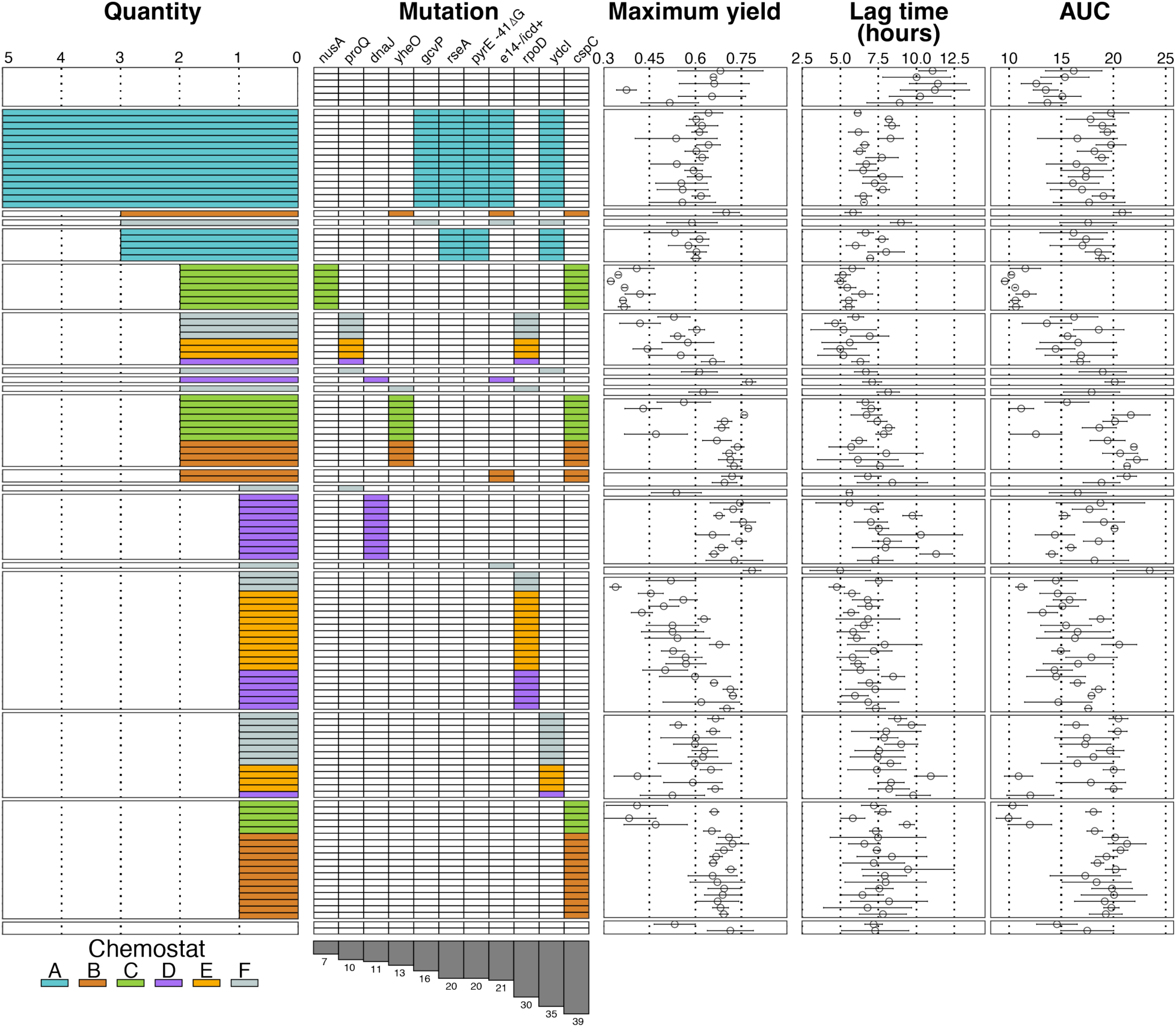
Other growth parameters extrapolated from growth curves. Other common metrics [maximum yield, lag time, and area under the curve (AUC)] calculated from growth curves measured in acetate in ancestor and evolved strains, organized by mutation as in Figure 5. Wild-type shown in the top panel. There was not a huge change in overall yield in response to our ALE, apart from *cspC* I20M/*nusA* R258C mutants (see Figure 6), which showed a decrease in overall yield. There was a significant decrease in overall lag time of ∼ 3 hours in the evolved clones relative to the wild-type (mean lag WT=10.48 hours, mean lag evolved=7.14 hours, *t*-test, *t*=10.86, n=524, *p*<0.0001). Finally, there was an increase in the AUC, a parameter that integrates different growth phases, in evolved strains relative to the wild-type which seems to correlate with maximum growth rate shown in Figure S7.

**Figure S11.**
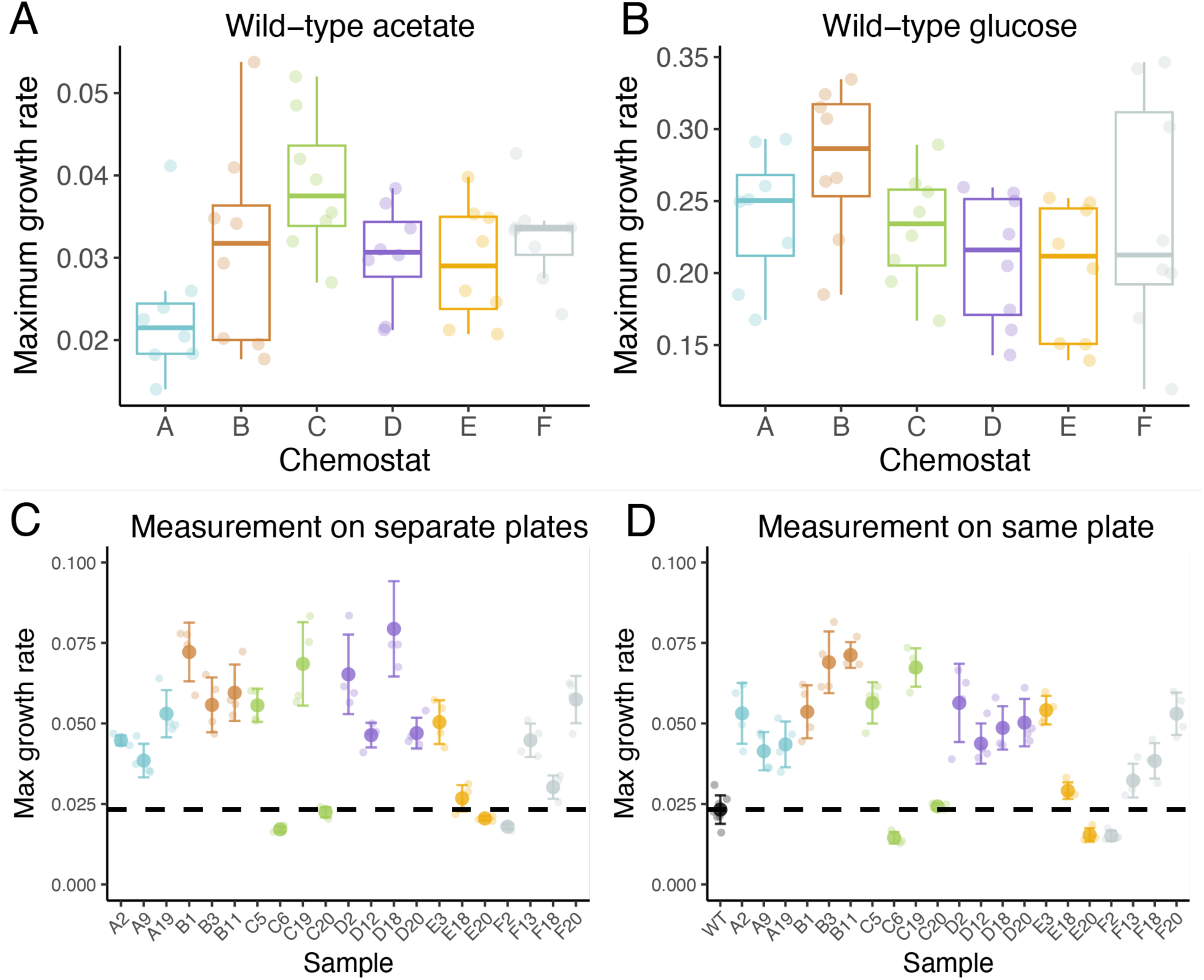
No evidence of significant plate effects during the measurement of maximum growth rate. To determine if measuring the maximum growth rate of isolated clones from different chemostats in separate plates resulted in plate effects, we first compared the wild-type comparison between plates for measurements in both acetate (A) and glucose (B). There was little significant difference between WT measurements for different plates (acetate one-way ANOVA, F_5,42_=3.02, *p*=0.020, pair-wise difference assessed with Tukey HSD; glucose: one-way ANOVA, F_5,42_=1.95, *p*=0.106, pair-wise difference assessed with Tukey HSD), suggesting minimal plate effects between different plate measurements. Furthermore, we also measured clones from different chemostats, originally measured on the different plate (C) on the same plate (D). Overall, there was not a huge difference when measured on different plates relative to within the same plate.

Table S1. Source data and statistics for growth rate of single-colony isolates.

Table S2. Excel file containing mutation information

